# Transretinal migration of astrocytes and brain/spinal cord-like cells arising from transplanted human retinal organoids

**DOI:** 10.1101/2022.05.12.491655

**Authors:** Ying V. Liu, Clayton P. Santiago, Akin Sogunro, Gregory J. Konar, Ming-wen Hu, Minda M. McNally, Yu-chen Lu, Zhuo-lin Li, Dzhalal Agakishiev, Sarah E. Hadyniak, Katarzyna A. Hussey, Tyler J. Creamer, Linda D. Orzolek, Derek Teng, Jiang Qian, Zheng Jiang, Robert J. Johnston, Seth Blackshaw, Mandeep S. Singh

## Abstract

Human retinal organoid transplantation can potentially restore vision in patients with degenerative retinal diseases. How the recipient retina regulates the maturation, fate specification, and migration of transplanted organoid cells is unknown. We transplanted human retinal organoid-derived cells into photoreceptor-deficient mice, conducted histology and single-cell RNA sequencing analyses, and observed two main classes of graft-derived cells. The first class consisted of retinal astrocytes and brain/spinal cord-like neural precursors, absent or rare in cultured organoids, that migrated into all recipient retinal layers and traveled long distances. The second class consisted of retinal progenitor-derived cells, including rods and cones, that remained in the subretinal space and matured more rapidly than photoreceptors in culture. These data suggest that the recipient subretinal space promotes the maturation of transplanted photoreceptors while inducing or expanding migratory cell populations that are not normally derived from retinal progenitors. These findings have important implications for cell-based treatment of retinal diseases.

## INTRODUCTION

Transplantation of immature retinal cells, such as photoreceptor precursor cells and retinal progenitor cells, has the potential to restore function to the degenerated or dysfunctional human retina. Vision loss may be retarded or reversed by direct cellular integration or cytoplasmic materials transfer ^1-13^. Stem cell-derived retinal organoids ^14-25^ are potential sources of renewable and standardized donor cells for therapy. Successful transplantation of these cells will require an understanding of the interactions between donor cells and the recipient environment. Here we address this challenge by transplanting human retinal organoid-derived donor cells subretinally into recipient mice and examining how the recipient environment impacts donor cells.

A substantial body of data in multiple animal models supports the regenerative potential of nonmigratory donor photoreceptor precursor-derived cells that mature in the recipient subretinal space, spurring experiments in large animals ^21, 26-30^ *en route* to clinical studies ^31-36^. Aside from the nonmigratory cells, migratory donor cells in the inner retinal layers overlying the graft have been observed ^37-46^, which do not appear to be essential for the therapeutic mechanism. These observations raised concerns regarding long-range migration of donor cells beyond the graft margins that may incite immune exposure and invasive tissue damage. In the few studies of photoreceptor transplantation in nonhuman primate recipients ^21, 29^, migratory cells have occasionally been observed ^29^, raising the possibility that it may also occur in humans. Studying the migration of donor cells may also illuminate strategies to regulate their spatial targeting, as with retinal ganglion cell homing in response to an induced chemokine gradient ^47^. Here, we assess the effects of the recipient retinal environment on the maturation and migratory behavior of transplanted cells derived from human retinal organoids. We molecularly define the fates of the migratory cells that leave and the nonmigratory photoreceptors that mature in the subretinal graft.

Whereas cell movement is not readily observed in the normal and pathological adult retina, retinal progenitor cells and their descendants undergo considerable radial migration and limited tangential migration during neurogenesis ^48-50^. In contrast, other CNS cells, including GABAergic neural precursors in the telencephalon ^51^ and optic nerve-derived astrocytes in the retina ^52^, undergo long-distance tangential migration. Similarly, stem cells and their progeny migrate following transplantation into the CNS ^53-56^. Thus, each developmental or transplantation context differentially regulates cell fate specification and migration.

The origin, heterogeneity, spatial distribution, and proliferative capacity of migratory donor cells have not been characterized. Migratory cells can arise from human retinal organoids ^29, 41, 43-46^ and human fetal retina ^42, 57^, suggesting that migratory behavior is not unique to donor cells obtained through pluripotent stem cell culture. Migratory donor cells were observed in mouse, rat, cat, and nonhuman primate retinas, and in normal and degenerative retinas ^29, 37-41, 58, 59^, suggesting that this phenomenon is common across recipient species and independent of retinal health. Few migratory cells were reported to express GFAP ^37, 58^, progenitor markers (e.g., PAX6, Nestin) ^40, 58^ or neuronal markers (e.g., MAP2, β-tublin3) ^60^, but not mature photoreceptor markers ^29, 39, 41, 43^-^46, 58, 61^. Gasparini *et al*., in a study of the influence of the murine retinal environment on the morphological maturation of transplanted donor human cone photoreceptors, also observed migratory cells but did not molecularly characterize these cells ^62^. Our studies use an unbiased approach to molecularly characterize the fate and maturation state of migratory and nonmigratory cells that arise from human retinal organoids upon transplantation into mice.

To understand how the recipient environment influences donor cells, we transplanted dissociated cells from human retinal organoids into mouse retinas and used imaging and single cell transcriptomics to characterize the fates, maturities, and migratory activities of graft-derived cells. We identified donor-derived migratory astrocytes and brain/spinal cord-like neural precursors that do not normally arise from retinal progenitor cells and nonmigratory photoreceptors that matured more rapidly in the subretinal space. Our findings highlight a key strength of organoid-derived cell transplantation in promoting photoreceptor maturation and a potential weakness in the expansion of a population of migratory astrocytes and brain/spinal cord-like neural precursor cells.

## RESULTS

### Human donor cells migrate from or remain in the subretinal space

To determine how the recipient subretinal space affects donor cells, we differentiated human retinal organoids, transplanted them into recipient mice, and later assessed donor cell position, fate, and maturity. To generate recipient mice, we crossed and bred mice with immune deficiency and retinal degeneration (supplementary **Fig. S1**). These C3H/HeJ*-Pde6b*^*Rd1/Rd1*^(*Rd1*) and NOD.Cg*-Prkdc*^*scid*^*/*J (*NOD/Scid*) double mutant mice are termed *Rd1/NS*. To generate donor cells, we used H9 human embryonic stem cells (hESCs) carrying a reporter that is expressed in all photoreceptors (CRX:tdTomato) ^63^. We used a gravity aggregation approach to differentiate stem cells into retinal organoids with robust generation of photoreceptors ^15, 22, 24^. On day 134 of organoid culture, we micro-dissected the human retinal organoids and transplanted the fragments into the subretinal space of recipient eyes (n=16 eyes). Four and a half months later, we evaluated the transplants.

As homozygosity for the *Rd1* allele causes virtually all photoreceptors to degenerate by adulthood in mice ^64^, distinct recipient outer nuclear and outer plexiform layers were not observed but the inner nuclear, inner plexiform, retinal ganglion cell (RGC), and retinal nerve fiber layers (collectively, the “inner retina”) were present.

We determined the positions of donor cells relative to the subretinal transplantation site. We identified all human donor cells based on immunolabeling for human nuclear antigen (HNA), or human ATP-dependent DNA helicase 2 subunit (Ku80 protein). We identified human donor photoreceptors based on transgenic expression of CRX:tdTomato (**Fig. 1A**). We observed two main classes of donor cells: (1) human cells in the recipient subretinal space (“nonmigratory cells”) that were photoreceptor or non-photoreceptor cells (**Fig. 1A)**, and (2) human cells in the recipient inner retina (“migratory cells”) that were not photoreceptors (**Fig. 1A**), suggesting that this population had migrated from the graft. Migratory cells were observed in the recipient inner retinal layers overlying the graft (“radial migration”, **Fig. 1B)**, whereas others had migrated away from and beyond the edges of the graft (“tangential migration”, **Fig. 1C**). Migratory cells traveled into all retinal layers, including the retinal ganglion cell (RGC) layer, inner plexiform layer (IPL), inner nuclear layer (INL), and retinal pigment epithelium/choroid (RPE/C) layer (**Fig. 1E)**. Of the tangential migratory cells (n=2,378 cells), 98.9% were within 1500 µm of the edges of the graft. The remaining 1.1% traveled beyond 1500 µm and were located exclusively in the RGC layer (**Fig. 1F**). We detected several migratory human cells in the regions flanking the optic nerve (“peripapillary migration”, **Fig. 1G)** but none in the optic nerve. We next sought to molecularly classify the fates of these nonmigratory and migratory cells.

**Fig. 1.**
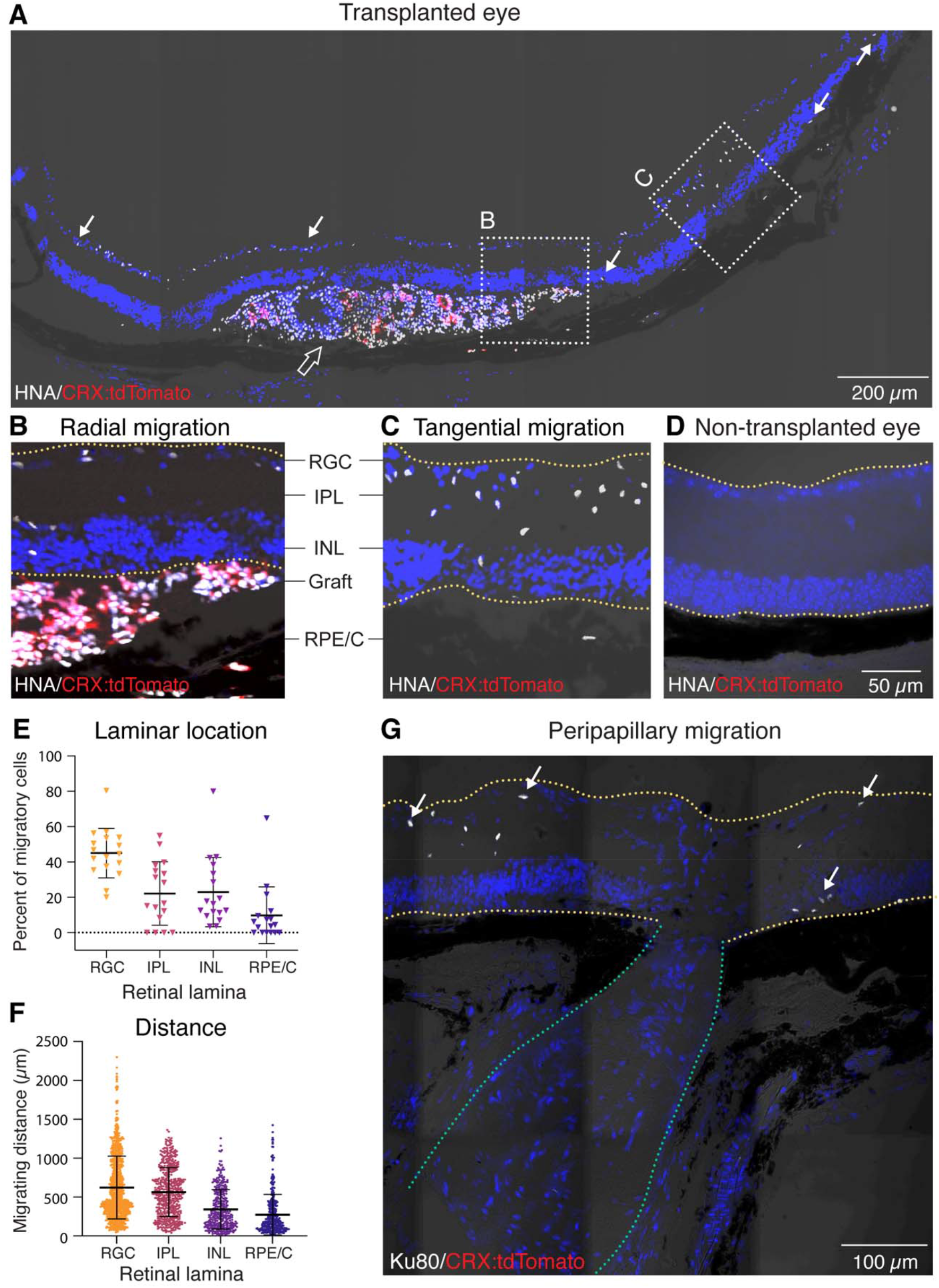
Human donor cells migrate from or remain in the subretinal space. **(A)** Immunohistochemical (IHC) staining of human nuclear antigen (HNA) showed migratory (arrows) and nonmigratory (empty arrow) cells of donor human retinal organoids in the recipient retina. Transplanted photoreceptors were identified by CRX:tdTomato reporter. **(B-C)** Migratory cells were detected overlying the graft (radial migration) and beyond the graft edge (tangential migration). **(D)** A control non-transplanted *Rd1/NS* mouse eye showed negative staining for HNA and CRX:tdTomato. **(E)** Relative abundance of migratory human cells in recipient retinal layers (RGC, IPL, INL, RPE/C) (n=17 sections from five transplanted eyes). **(F)** Quantification of the migrating distance of the human cell nuclei from the graft edge in different retinal laminae (RGC, IPL, INL, RPE/C) (n = 21 sections from five transplanted eyes). **(G)** Migratory donor human cells (identified using Ku80) were detected in the regions flanking the optic nerve, i.e., peripapillary migration. Yellow lines in **B, C, D**, and **G** delineate the boundaries of the recipient retina. Green lines in **G** delineate the optic nerve. *Abbreviations: RGC: retinal ganglion cell; IPL: inner plexiform layer; INL: inner nuclear layer; RPE/C: retinal pigment epithelium and choroid*.

### Donor cells adopt retinal-derived and non-retinal-derived cell fates

To determine how the recipient subretinal microenvironment affects the gene expression and cell fate specification of the migratory and non-migratory donor cells, we conducted single cell RNA sequencing on cells from human retinal organoids transplanted and matured *in vivo* (“transplanted organoids”) and from age-matched organoids that were maintained *in vitro* (“cultured organoids”) (**Fig. 2A**). We analyzed a total of 5,831 human cells that were recovered from the transplanted (1,561 cells) and cultured (4,270 cells) organoids. We identified retinal cell types including retinal progenitor cells (RPCs), photoreceptor precursor cells, rods, cones, bipolar cells, horizontal cells, and Müller glia based on their gene expression profiles (**Fig. 2B-D**). The quantities of cones, bipolar cells, and horizontal cells were similar in the transplanted and cultured organoids. In contrast, retinal progenitor cells (RPCs), photoreceptor (PR) precursor cells, and Müller glia were more abundant in cultured organoids, whereas rods were more abundant in transplanted organoids (**Fig. 2E**). The smaller populations of RPCs and photoreceptor precursor cells and larger population of rods in the transplanted organoids suggest that the recipient microenvironment promotes specification and maturation of retinal cell fates.

**Fig. 2.**
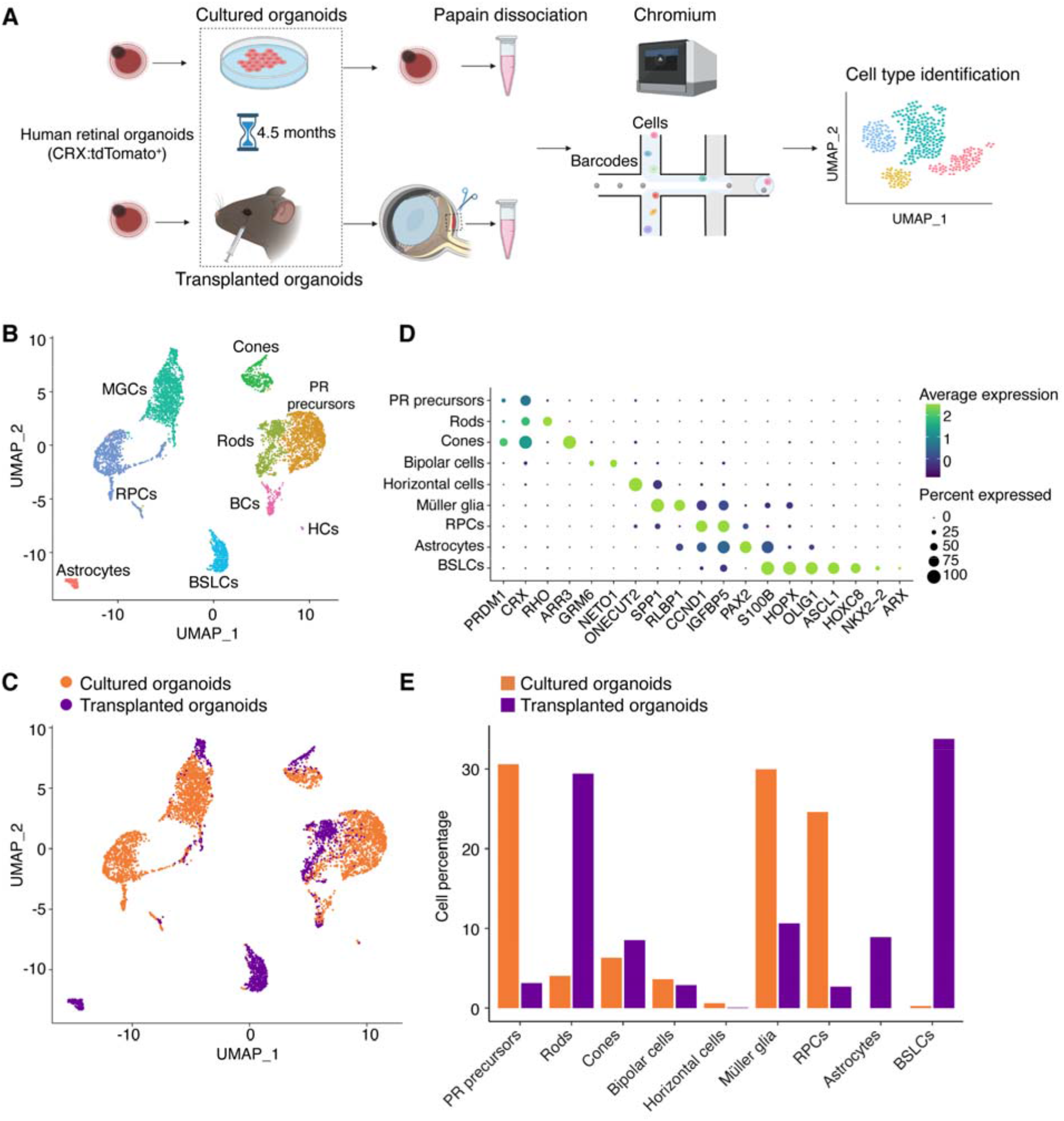
Single-cell RNA sequencing (scRNA-seq) analysis identified retinal-derived and non-retinal-derived cell fates in transplanted and cultured retinal organoid cells. **(A)** Schematic showing the *in vivo* and *in vitro* conditions of the donor cells ultimately analyzed by scRNA-seq. CRX:tdTomato^+^ hESC derived-retinal organoids (aged 134 days) were transplanted into *Rd1/NS* mice or maintained in culture. Four and a half months later, single cell suspensions of transplanted and cultured retinal organoids (age-matched) were collected by papain dissociation and analyzed by Chromium scRNA-seq. **(B-C)** scRNA-seq identified nine transcriptionally distinct cell clusters from the pool of transplanted and cultured retinal organoid cells (n=5,831 cells). **(D)** Dot plots of marker gene expression in the identified cell clusters. The color scale corresponds to the average gene expression and the dot size corresponds to the percent of positively expressing cells in each cluster. **(E)** The relative abundance of cells of each type in transplanted and cultured retinal organoids.

In addition to these cell types, we identified two cell clusters that could not be ascribed solely to known retinal-derived cell fates. The cells in one cluster expressed genes that are broadly expressed in retinal and other CNS progenitors such as *ASCL1* (**Fig. 2D**) and *HES6* (supplementary **Fig. S2**). They also expressed genes that are not normally detected in the developing retina including *NKX2*-2 and *ARX*, both of which are prominently expressed in ventral telencephalic and diencephalic neural progenitors, as well as *HOXC8*, whose expression is normally restricted to the developing spinal cord (**Fig. 2D**). Based on this gene expression profile, we designated the cells in this cluster as “brain and spinal cord-like” (BSL) cells. BSL cells comprised approximately 1% of cells in the cultured organoids but were over 30 times more abundant in the transplanted organoids (**Fig. 2E**). Cells in the second cluster expressed markers characteristic of retinal astrocytes, such as *PAX2* and *S100B* (**Fig. 2D**). Normally, retinal astrocytes are born in the optic nerve head and migrate into the retina. Strikingly, astrocytes were entirely absent in the cultured organoids, but comprised approximately 8% of cells in the transplanted organoids (**Fig. 2E)**. These data suggest that the recipient microenvironment directs some donor cells to assume fates that are not normally acquired by retinal progenitors.

### Migratory donor retinal astrocytes and brain/spinal cord-like neural precursors

Next, we connected the cell types identified by scRNA-seq analysis to their migratory or non-migratory properties. We computationally aggregated and scored the expression of genes associated with cell motility and migration by gene ontology (GO) classification (supplementary **data file S1**). We found that astrocytes and BSL cells had the highest average migration scores (**Fig. 3A**), suggesting that they were migratory cells.

**Fig. 3.**
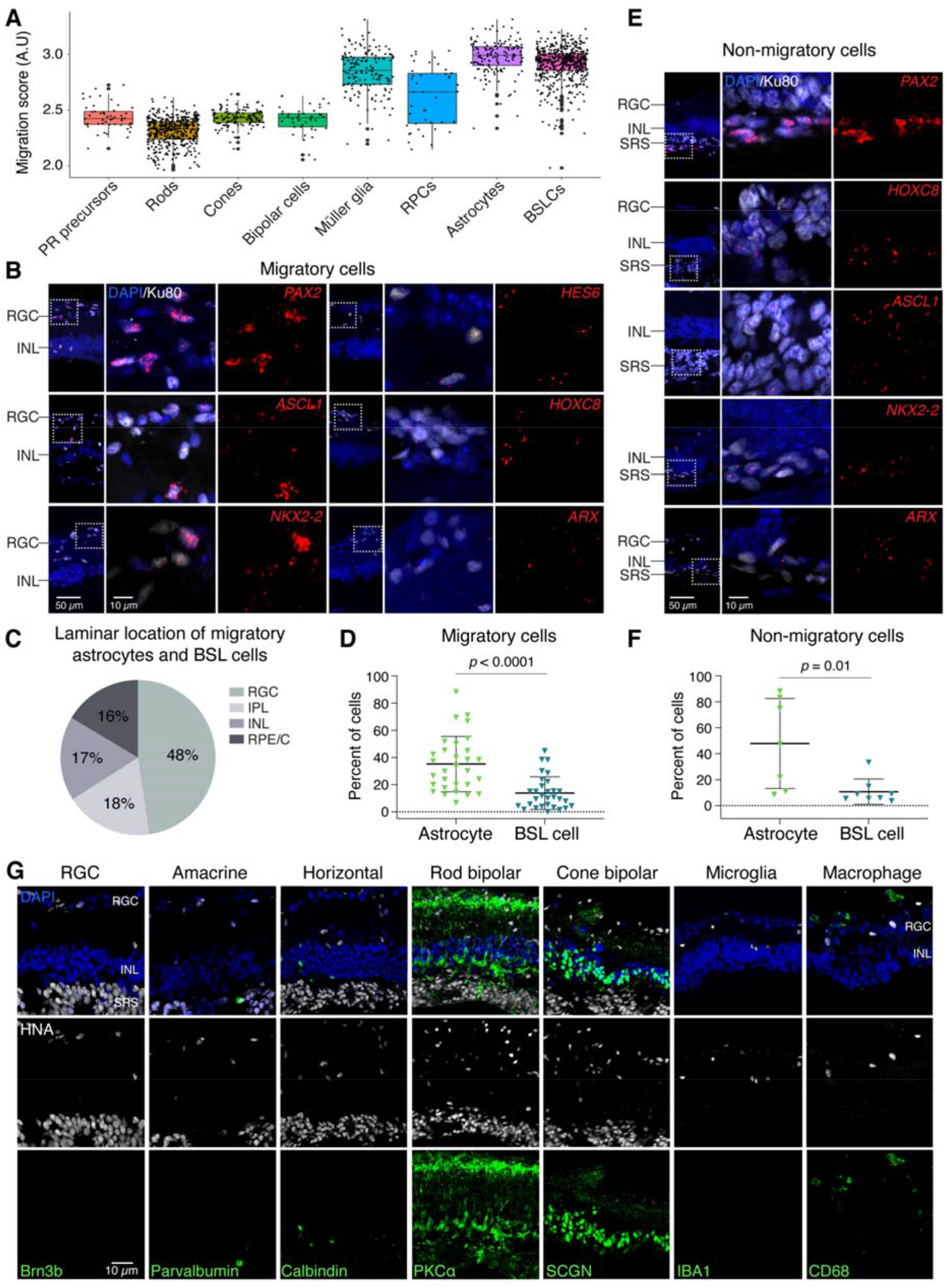
Donor retinal astrocytes and brain/spinal cord-like (BSL) neural precursors show migratory capacity. **(A)** Astrocytes and BSL cells show the highest migration score among the cell types identified in transplanted retinal organoids. **(B)** RNAscope staining showed migratory cells expressing markers of astrocytes (*PAX2, HES6*) and BSL cells (*ASCL1, HOXC8, NKX2-2, ARX*). IHC counterstaining of human nuclear antibody Ku80 was used to detect transplanted human cells. **(C)** Relative abundance of migratory astrocytes and BSL cells in different recipients’ retinal laminae (RGC, IPL, INL, RPE/C) by histological quantification. **(D)** Histological quantification of migratory astrocytes and BSL cells in migratory human cells (n=30 regions of interest from three to four transplanted eyes). **(E-F)** RNAscope staining (E) and quantification (F) of non-migratory astrocytes and BSL cells in the subretinal space (SRS) (n = 7-8 regions of interest from four transplanted eyes). **(G)** Migratory cells negatively expressed markers of RGC (Brn3b, human specific antibody), amacrine cells (parvalbumin, human specific antibody), horizontal cells (calbindin), rod bipolar cells (PKCa), cone bipolar cells (SCGN), microglia (IBA1), and macrophage (CD68). DAPI staining was performed to identify the nuclei of recipient retinal laminae. Anti-human nuclei antibody (HNA) was used to track transplanted human cells.

To test this hypothesis, we examined expression of genes including *PAX2, HES6, ASCL1, HOXC8, NKX2-2*, and *ARX*, that together identify astrocytes and BSL cells. We observed expression of these genes in migratory cells located in all recipient layers (**Fig. 3B-C**), suggesting that astrocytes and BSL cells had migratory capacity. Among migratory cells, human astrocytes were more abundant than BSL cells (**Fig. 3D**). Among non-migratory cells that remained in the subretinal space within the graft, we also observed astrocyte and BSL cells (**Fig. 3E-F**), suggesting that expression of astrocyte or BSL cell related genes alone was insufficient for migration.

Migratory cells were almost exclusively CRX:tdTomato^−^, consistent with these cells taking on astrocyte or BSL fate and not photoreceptor fate (see **Fig. 1A**). Rare CRX:tdTomato^+^ cells were detected in the RPE/C layer and were possibly misplaced photoreceptors (see **Fig. 1A**). Human migratory cells did not express established markers of RGCs, amacrine cells, horizontal cells, rod and cone bipolar cells, microglia, or macrophages (**Fig. 3G**, supplementary **Fig. S2**).

Together, these data suggest that some but not all donor human astrocytes and BSL cells were migratory cells whereas donor photoreceptors and other retinal neurons were non-migratory.

### Actively proliferating cells are rare among migratory and non-migratory donor cells

Migratory cells, especially if they are proliferative, may negatively impact the recipient. To determine the influence of the recipient microenvironment on the proliferation of migratory and non-migratory donor cells, we examined expression of the proliferation marker protein Ki-67. As expected, expression of Ki-67 was rarely observed in CRX:tdTomato^+^ photoreceptor precursors in cultured organoids, and were significantly less abundant in transplanted organoids (**Fig. 4A**). The smaller population of proliferating CRX:tdTomato^+^ cells in the transplanted organoids suggest that the *in vivo* microenvironment promotes maturation of postmitotic photoreceptor cells.

**Fig. 4.**
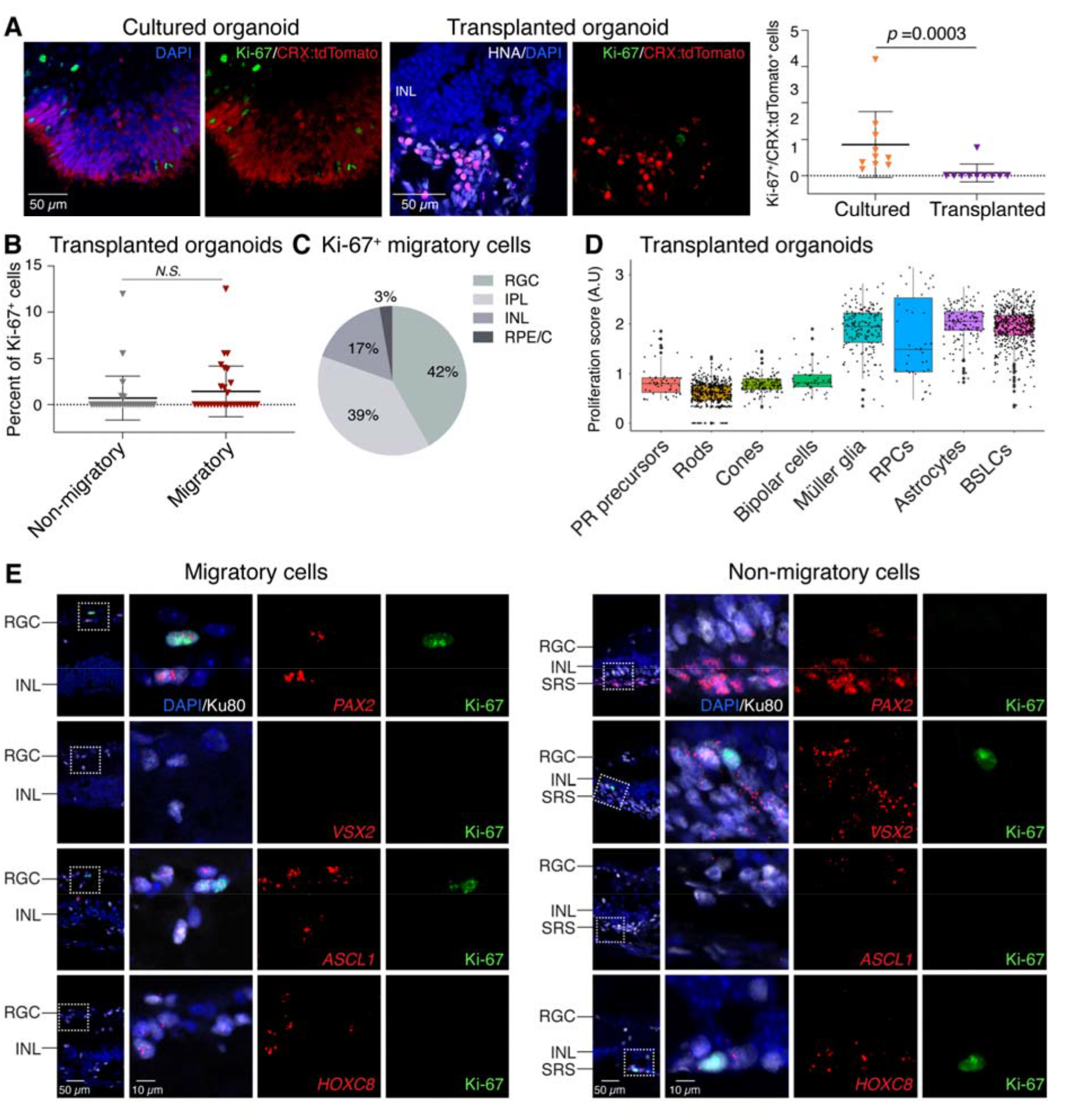
Actively proliferating cells are rare among migratory and non-migratory donor cells. **(A)** IHC staining showed that Ki-67^+^ proliferating cells were rarely observed in CRX:tdTomato^+^ photoreceptor precursors in cultured organoids, and were significantly less abundant in transplanted organoids (n = 10 sections from three individual samples per group). **(B)** Similarly, Ki-67^+^ cells were rarely detected among non-migratory and migratory cells in transplanted retinal organoids (n = 30 sections from five transplanted eyes). **(C)** Relative abundance of migratory Ki-67^+^ cells in different retinal laminae (RGC, IPL, INL, RPE/C). **(D)** ScRNA-seq analysis showed the highest proliferation score in Müller glia, retinal progenitor cells (RPCs), astrocytes, and BSL cells among the identified cell types in transplanted retinal organoids. **(E)** RNAscope and IHC counterstaining showed sparse Ki67^+^ cells in migratory astrocytes (*PAX2*^+^) and BSL cells (*ASCL1*^+^), as well as in non-migratory RPCs (*VSX2*^+^) and BSL cells (*HOXC8*^+^).

In eyes with transplanted organoids, 0.7% of non-migratory cells and 1.4% of migratory cells expressed Ki-67 **(Fig. 4B)**, and the difference between these values was not statistically significant. We observed that the few Ki-67^+^ migratory cells occupied all retinal laminae of the recipient (**Fig. 4C**).

To identify the proliferating cells, we developed a proliferation scoring system by computationally aggregating the expression level of proliferation-associated genes (supplementary **data file S2**). We found that astrocytes, Müller glia, RPCs, and BSL cells showed the highest proliferation score (**Fig. 4D**), suggesting that these cells were proliferating. To test this hypothesis, we examined expression of Ki-67 in *PAX2*^*+*^ (astrocytes), *VSX2*^*+*^ (RPCs), and *ASCL1*^*+*^ and *HOXC8*^*+*^ (BSL) cells. Precise quantification was impractical due to the rarity of double-positive cells. Nevertheless, we found a few migratory *PAX2*^+^ astrocytes, and very few migratory *ASCL1*^*+*^ BSL cells, that were Ki-67^+^. Proliferating Ki-67^+^/*VSX2*^*+*^ RPCs remained in the subretinal space (**Fig. 4E**).

Taken together, these data suggest that migratory proliferating donor human cells are rare and are mostly astrocytes, and that nonmigratory proliferating cells are rare and are mostly RPCs.

### Donor cones and rods mature more rapidly in the recipient subretinal space than in culture

Our scRNA-seq analysis suggested that the recipient subretinal space promotes photoreceptor fate and possibly maturation (see **Fig. 2E**). To test this hypothesis, we first assessed cone maturation. We evaluated the gene expression profiles of cones from transplanted organoids and cultured organoids using pseudotime analysis, comparing these cells to published datasets of embryonic, postnatal, and adult cones isolated directly from human retina ^65^. The transcriptional profiles suggested that the cones from transplanted organoids resembled adult cones, whereas the cones from cultured organoids more closely resembled embryonic cones (**Fig. 5A-B**). Expression of mature cone-specific genes were consistently higher in transplanted than in cultured cones (supplementary **Fig. S3**), including all three cone opsins (*OPN1LW, OPN1MW*, and *OPN1SW*) (**Fig. 5C**). The proportions of CRX:tdTomato^+^ cells that expressed L/M-opsin or S-opsin were significantly higher in transplanted organoids (L/M-opsin^+^: 26.4%, S-opsin^+^: 13.7%) compared to cultured organoids (L/M-opsin^+^: 2.7%, S-opsin^+^: 1.3%) (**Fig. 5D**). Similarly, the fraction of L/M-opsin^+^ or S-opsin cells^+^ with inner or outer segments (segment^+^) was significantly higher in transplanted organoids than cultured organoids (**Fig. 5E**). We measured the intrinsic electrical properties of a transplanted human cone cell and found large capacitance currents (∼2 nA), indicating relatively large cell membrane areas as normally observed in mature cones (**Fig. 5F**).

**Fig. 5.**
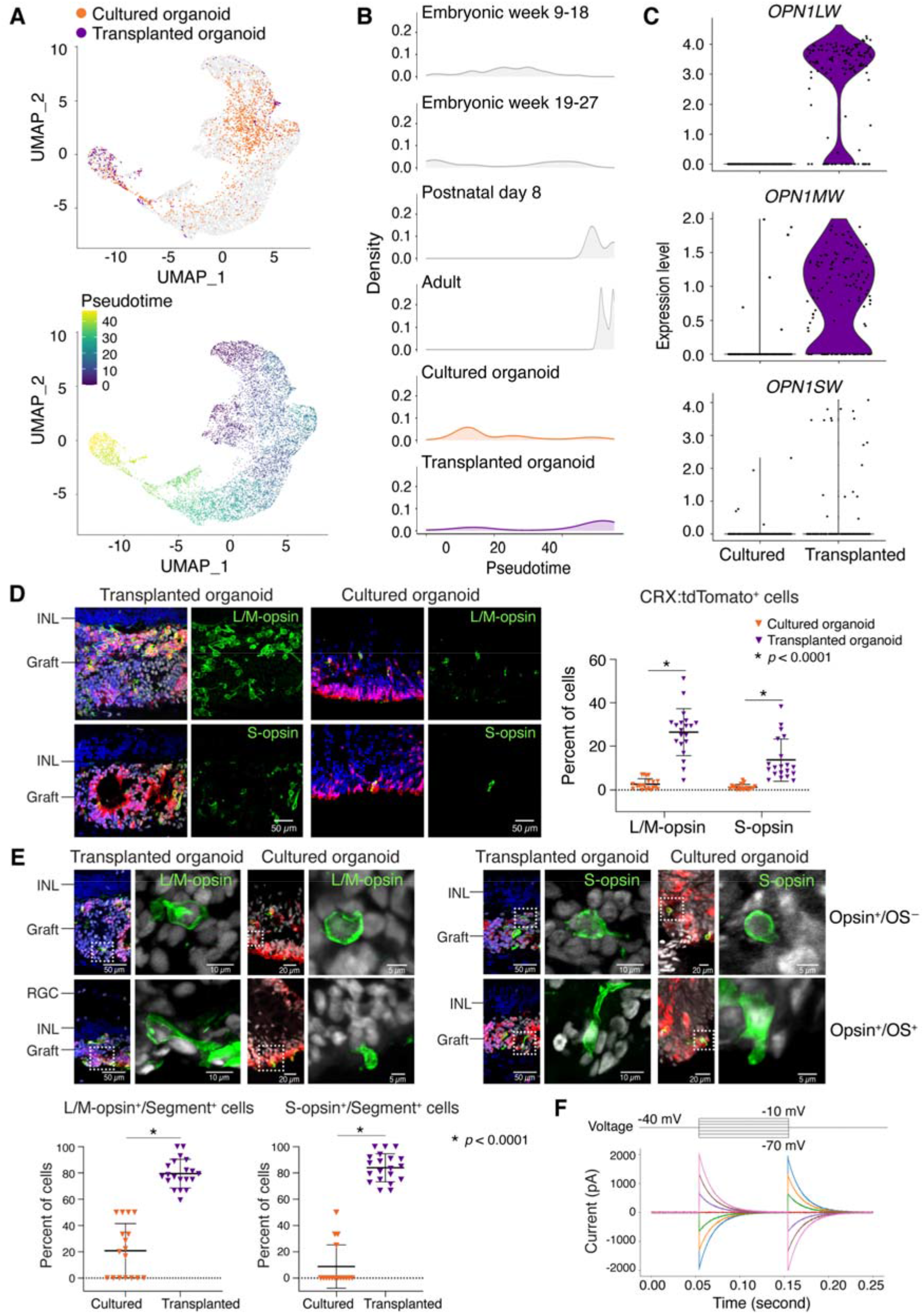
Donor cone photoreceptors mature more rapidly in the recipient subretinal space than in culture. **(A)** UMAP plots embedded the pseudotime maturation trajectories of cone photoreceptors in transplanted (n=3) and cultured retinal organoids (age-matched, n=2), comparing to normal human cone photoreceptor development (aged from embryonic week nine to adulthood). Cells are colored by cell type (top UMAP) and pseudotime (bottom UMAP). **(B)** Ridge plots showed that the transcriptional maturation of transplanted cone photoreceptors resembled adult human cone photoreceptors, whereas cultured cone photoreceptors resembled embryonic human cone photoreceptors. **(C)** scRNA-seq violin plots showed significantly more *OPN1LW, OPN1MW*, and *OPN1SW* expression in transplanted than cultured retinal organoids. **(D)** IHC staining and quantification showed significantly higher proportion of L/M-opsin^+^ or S-opsin^+^ photoreceptors (CRX:tdTomato^+^) in transplanted than cultured retinal organoids (n = 20 sections from four individual samples per group). **(E)** IHC images showed representative L/M-opsin^+^ or S-opsin^+^ cone photoreceptors with (OS^+^) or without (OS^-^) outer segments. Histological quantification showed the fraction of L/M-opsin^+^ or S-opsin^+^ cells with inner/outer segment formation (segment^+^) was significantly higher in transplanted than cultured retinal organoids (n=16-20 sections from four individual samples per group). **(F)** Single-cell patch-clamp recording of a transplanted human cone photoreceptor showed the mature cone-specific large-capacitance current.

Next, we evaluated rod maturation in transplanted organoids and cultured organoids. As with cones, gene expression and pseudotime analysis suggested that rods from transplanted organoids resembled adult rods, whereas rods from cultured organoids resembled embryonic rods (**Fig. 6A-B**). Expression of *RHO* (**Fig. 6C**) and other rod-specific genes (supplementary **Fig. S3**) was higher in rods from transplanted organoids than cultured organoids. The proportions of CRX:tdTomato^+^ cells that expressed Rho were significantly higher in transplanted organoids (61.5%) compared to cultured organoids (45.5%) (**Fig. 6D**). Similarly, the fraction of Rho^+^ cells with inner or outer segments (segment^+^) was significantly higher in transplanted organoids (87.4%) than cultured organoids (29.8%) (**Fig. 6E**).

**Fig. 6.**
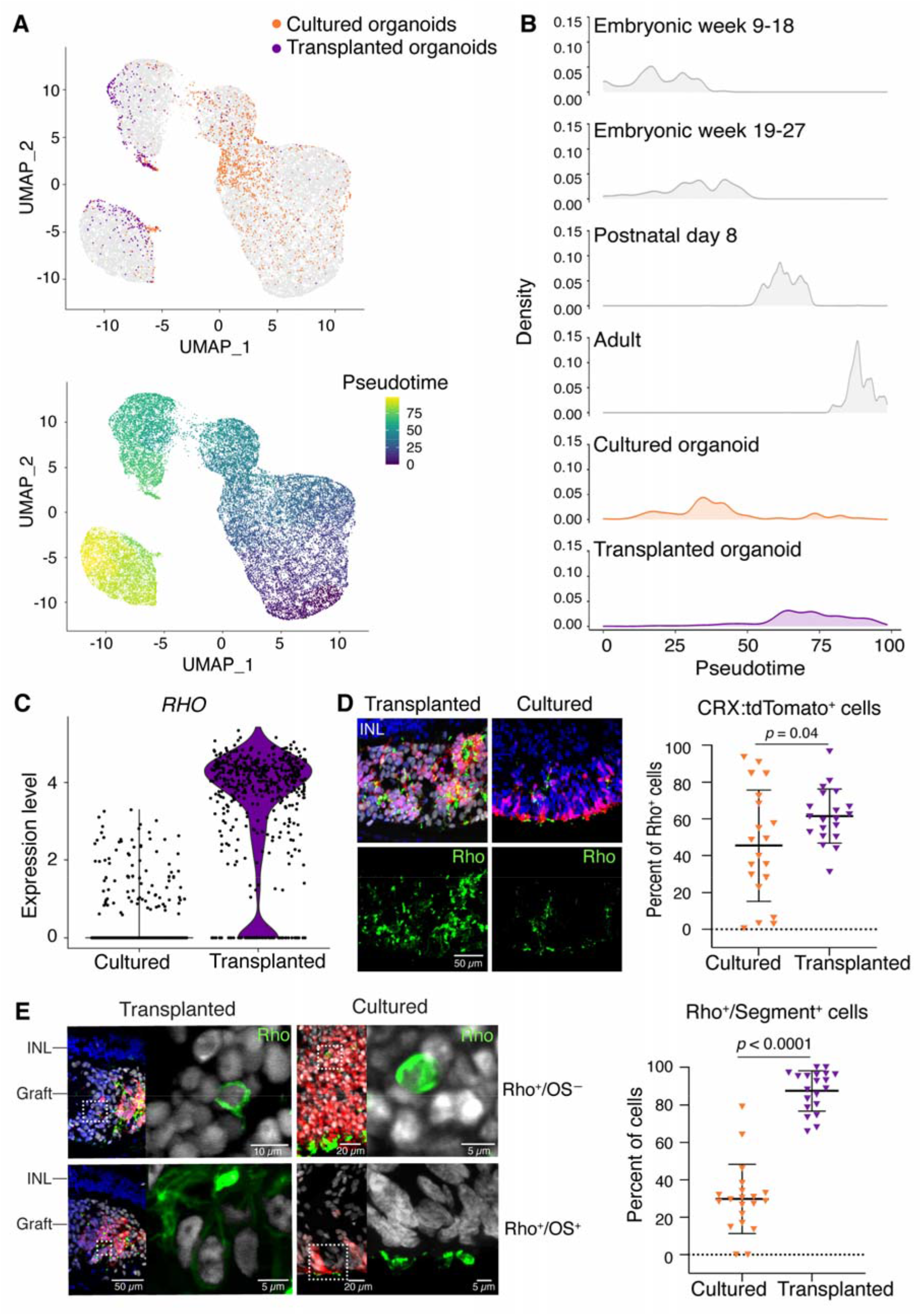
Donor rod photoreceptors mature more rapidly in the recipient subretinal space than in culture. **(A)** UMAP plots embedded the pseudotime maturation trajectories of rod photoreceptors in transplanted (n = 3) and age-matched cultured retinal organoids (n = 2), comparing to human rod development (aged from embryonic week nine to adult). Cells were colored by cell type (top UMAP) and pseudotime (bottom UMAP). **(B)** Ridgeline plot of scRNA-seq data indicated that the transcriptional maturation of transplanted rod photoreceptors resembled adult human rod photoreceptors, whereas cultured rod photoreceptors transcriptionally resembled embryonic human rod photoreceptors. **(C)** Violin plots of scRNA-seq analysis showed upregulation of *RHO* gene expression in transplanted retinal organoids compared to cultured retinal organoids. **(D)** IHC staining and quantification showed significantly higher proportion of CRX:tdTomato^+^ photoreceptors that expressed Rho^+^ in transplanted than cultured retinal organoids (n = 20 sections from four individual samples per group). **(E)** Representative IHC images showed Rho^+^ rod photoreceptors with (OS^+^) or without (OS^-^) outer segments. Histological quantification showed a significantly greater fraction of Rho^+^ cells with inner/outer segment formation (segment^+^) in transplanted than cultured retinal organoids (n=20 sections from four individual samples per group).

Finally, we investigated general features of photoreceptor maturity. Expression of certain synaptic proteins was upregulated in cones and rods in transplanted organoids compared to cultured organoids (supplementary **Fig. S4 A**). In CRX:tdTomato^+^ donor photoreceptors, the number of CtBP2^+^ puncta, which marks synaptic ribbons, was significantly higher in cells from transplanted organoids compared to cultured retinal organoids (supplementary **Fig. S4 B**).

Together, these data suggest that the recipient subretinal space, compared to the *in vitro* environment of cultured organoids, promotes maturation of rods and cones.

## DISCUSSION

In these studies, we observed two major differences between cells from donor retinal organoids transplanted into mice and cells from chronologically equivalent retinal organoids maintained in culture. The transplanted cells were maintained in the degenerative recipient subretinal space for several months, therefore simulating conditions directly relevant for cell-based therapies for photoreceptor dystrophy. The most prominent and unexpected difference was the observation of migratory donor astrocytes and BSL cells in the transplanted cell population. Astrocytes and BSL cells underwent radial migration into, and long-distance tangential migration along, all retinal laminae (apart from the outer nuclear layer of photoreceptor cells that was absent in the degenerate recipients). The migratory astrocytes and BSL cells were generally non-proliferative, although graft-derived retinal progenitors showed proliferation without migration. In contrast to these migratory cells, transplanted photoreceptors, inner retinal neurons, and Müller glia were non-migratory and remained in the subretinal transplant site. The second major difference between transplanted and cultured organoids pertained to photoreceptor maturity. Based on gene expression and morphology, transplanted rods and cones were more mature than photoreceptors from cultured organoids. These data expand our understanding of photoreceptor and non-photoreceptor development in transplanted retinal organoids and highlight the importance of unbiased approaches to cell fate identification and spatial tracking following organoid transplantation.

The migratory astrocytes and BSL cells from transplanted organoids display molecular profiles distinct from cells in mature cultured organoids. The astrocytes express *PAX2*, which normally delineates the optic stalk *in vivo* ^66^. *PAX2* is detected in retinal progenitors in early-stage retinal organoids but is undetectable at later stages ^65, 67^. Moreover, cultured retinal organoids have not been reported to generate astrocytes *in vitro*. The BSL cells express *ASCL1, HOXC8, NKX2-2*, and *ARX. NKX2-2* and *ARX*-expressing cells are found in very early-stage retinal organoids, but not after 60 days in culture ^65^. *HOXC8* expression is normally restricted to the posterior spinal cord and is absent from developing human retina and retinal organoids ^65, 67^. Though *PAX2*^+^ astrocytes and *ARX*^+^ telencephalic interneurons undergo long-distance tangential migration *in vivo* ^*51, 52*^, astrocyte or BSL identity was not sufficient to induce migration of graft-derived cells, as many astrocytes and BSL cells remained localized in the subretinal space. Our experiments lacked the temporal resolution to determine whether transplantation induced trans-differentiation of cells that initially adopted retinal identity or selectively promoted the proliferation of small numbers of residual BSL cells.

Prior publications have shown migratory transplanted cells, but their capacity to proliferate and migrate long distances were not known. Seiler and colleagues noted migratory human donor cells six months after transplantation of early-stage hESC-derived retinal organoids in the retinal degeneration nude rat ^41^. Using LMNB2 to identify human donor cells, Lamba and colleagues reported occasional migratory human induced pluripotent stem cell (iPSC)-derived PAX6^+^ and GFAP^+^ cells at 2 months ^58^. In another study, migratory cells were seen just seven days after subretinal delivery of human fetal CD29^+^/SSEA1^+^ donor cells ^68^, suggesting that migration occurs soon after transplantation. Whether or not the early-migratory cells are the same as those observed months later is unknown. In the wild-type cat ^59^, enhanced immunosuppression appeared to lead to greater cell migration, suggesting a role of immune cells. It is not known whether migratory donor cells negatively affect recipient retinal function or if depletion is required prior to transplantation.

The cues in the recipient environment that promote migratory cell fate and maturation of photoreceptors are not known. Multiple cell-extrinsic cues regulate cell specification in human retinal organoids. Dynamic regulation of thyroid hormone and retinoic acid signaling specifies cone subtypes in human retinal organoids ^24, 69^. Though the roles of these cues in the subretinal environment following transplantation is not understood, they potentially regulate photoreceptor specification and maturation.

In conclusion, we found that the murine recipient subretinal environment affects human stem cell derived retinal organoid cells in two distinct ways. First, the recipient environment promotes a population of organoid-derived astrocytes that are capable of radial and tangential migration. Second, the recipient environment promotes the maturation of organoid-derived rod and cone photoreceptors that remain in the subretinal space. These results may inform future research on the consequences of donor cell migration in transplant recipients, methods to purify retinal organoid-derived cells, and pharmacological strategies to accelerate the maturation of donor photoreceptor cells. A deeper understanding of these issues may help guide the development of safe and effective clinical treatment involving the replacement or augmentation of human stem cell derived retinal photoreceptor precursors for the purposes of restoring visual function.

## METHODS

### Cell culture and retinal organoids differentiation

The use of human stem cells was approved by the Johns Hopkins ISCRO (ISCRO00000249). The CRX:tdTomato H9 human embryonic stem cell line (hESCs) ^63^ was cultured following the gravity aggregation approach to differentiate retinal organoids, as previously described ^15, 22, 24^. Media used for retinal organoids culturing were listed in supplementary **Table S1** and **Table S2**. On day 134, retinal organoids were used for transplantation. Detailed protocol was described in Supplementary Materials and Methods.

### Recipient mice

All animal experiments were carried out in accordance with the ARVO Statement for the Use of Animals in Ophthalmic and Vision Research. All procedures were approved by the Johns Hopkins University Animal Care and Use Committee (approval M016M17). We created a recipient mouse model with immune-deficiency and retinal degeneration (referred to as *Rd1/NS*) by crossbreeding *Rd1* mice and *NOD/Scid* mice. Mice were genotyped by Transnetyx Tag Center (Cordova,TN, USA) (see primers in supplementary **Table S3**), and characterized by immunohistochemistry (IHC) staining and flow cytometry analysis. Detailed protocol was provided in Supplementary Materials and Methods.

### Transplantation

Donor retinal organoid cells (harvested as micro-dissected multilayered retinal fragments) were obtained from CRX:tdTomato^+^ hESC-derived retinal organoids (aged 134 days, n=4). The isolated retinal fragments were transplanted into the subretinal space of *Rd1/NS* mice (aged 6 to 8 weeks, n=16 eyes). Detailed protocol was described in Supplementary Materials and Methods.

### Single cell RNA sequencing

Four and a half months post-transplantation, single cell RNA sequencing (scRNA-seq) was performed on dissociated single cells from transplanted (n=3 eyes) and cultured retinal organoids (n=2) (age-matched) using the Chromium platform (10X Genomics). Quality control of scRNA-seq data were shown in supplementary **Fig. S5**. Detailed protocol and data processing were provided in Supplementary Materials and Methods.

### Histological analysis

Four and a half months post-transplantation, the recipient mice eyes and cultured retinal organoids were fixed with 4% paraformaldehyde (PFA) (Electron Microscopy Sciences, Hatfield, PA, USA) in PBS and dehydrated in a sucrose gradient (10%, 20%, 30%), then blocked in optimal cutting temperature compound (OCT) (Sakura Finetek, Torrance, CA, USA). Seven to ten micrometer sections of recipient eyes and cultured organoids were used for RNAscope and IHC counterstaining. RNAscope and IHC counter-staining was performed according to the manufacturer’s protocol (Advanced Cell Diagnostics (ACD), see Protocol #MK 51-150, Appendix D.). The RNA probes, fluorophores, primary antibodies, and secondary antibodies used were listed in supplementary **Table S4**. Negative and positive multiplex control probes staining were run in parallel with the target probes following the same protocol (data shown in supplementary **Fig. S6**). The primary antibodies and secondary antibodies used for IHC staining were listed in supplementary **Table S5**. Detailed protocol and histological quantification were presented in Supplementary Materials and Methods.

### Electrophysiology

The electrophysiological recording was performed on the transplanted photoreceptors eight months post-transplantation to measure their physiological properties. Retinas with transplanted retinal organoids were dissected under infrared light and sectioned into 200µm slices, then transferred to a recording chamber. The CRX:tdTomato^+^ photoreceptors of the transplanted retinal organoids were targeted under an epifluorescence microscope for consequent whole-cell patch-clamp recording (see details in Supplementary Materials and Methods).

### Statistical analysis

Quantitative histology data were analyzed using two-way ANOVA. Sidak’s test or Tukey’s test was adopted for multiple comparisons (two-tailed). Independent T-test or Mann-Whitney U test was used for two variants comparison. Statistical analysis was carried out using SPSS software (version 25, IL, USA). *p* < 0.05 was taken to be significant. Statistical data were presented as mean ± SD. Graphs were drawn with GraphPad Prism software (version 8, CA, USA). Schematics were created with BioRender.com (agreement number QH23QWJX12, KY23QWKEPB).

## Data availability

The raw scRNA-seq data and count matrices generated during this study can be accessed at GEO accession number GSE197847. The merged count matrices and cell metadata are also available for download at https://github.com/csanti88/transplant_human_organoid_retina_2022. Interactive queries of individual gene expression patterns can be performed on the UCSC cell browser at https://xeno-hesc-retina.cells.ucsc.edu^*70*^.

## Acknowledgements

This work was funded by the following funding: NEI R01EY033103 (MSS), Foundation Fighting Blindness (MSS), Stein Innovation Award from Research to Prevent Blindness (SB), the Shulsky Foundation (MSS), the Joseph Albert Hekimian Fund (MSS), the Juliette RP Vision Foundation (YL), Research to Prevent Blindness (unrestricted grant to the Wilmer Eye Institute at Johns Hopkins University and the Cullen Eye Institute at Baylor College of Medicine), NEI Core Grant EY001765, Visual Sciences Training grant 2T32EY007143 (CPS), NSF DGE-1746891 (KH), NIH F31EY029157 (SEH), NEI R01EY030872 (RJJ). We thank Rhonda Griebe and Mary Ellen Pease (Wilmer Microscopy and Imaging Core Facility) for their kind assistance. We thank Dr. Kang V. Li for his technical assistance, and Wendy Yap for comments on the manuscript. We would also like to thank Brittney Wick, Matthew Speir and Maximilian Haeussler with their assistance with setting up and hosting the data on the UCSC cell browser.

## Author contributions

MSS, SB, RJJ conceived the experiments. YVL, CPS, AS, GJK, SEH, KAH, TJC, LDO, ZJ performed the experiments. All the authors contributed to data analysis and approved the final manuscript.

