## Supplemental text and figures S1-S6 for "Transretinal migration of astrocytes and brain/spinal cord-like cells arising from transplanted human retinal organoids"

Ying V. Liu, *et al*.

### Supplementary Materials

#### Supplementary Materials and Methods

##### *Cell Lines*

The use of human stem cells was approved by the Johns Hopkins ISCRO (ISCRO00000249). The H9 CRX:tdTomato human embryonic stem cell line (hESCs) was a kind gift from Dr. David M. Gamm (University of Wisconsin Hospitals, USA). Stem cells were maintained in mTeSR1 (Stem Cell Technologies, Cambridge, MA, USA) on 1% (vol/vol) Matrigel-GFRTM (BD Biosciences, USA, No. 354230,) coated dishes and grown in a 37°C HERAcell 150i incubator at 10%$CO_2$ and 5% $O_2$ incubator (Thermo Fisher Scientific, MA, USA). Cells were passaged upon confluence (every 3-6 days) using Accutase (Sigma-Aldrich, MO, USA, No. SCR005) for 7–10 minutes, and dissociated to single cells. Cells in Accutase were added 1:2 to mTeSR1 plus 5 µM Blebbistatin (Bleb; B0560, Sigma), pelleted at 700 g for 5 minutes, and suspended in mTeSR1 plus Bleb and plated at 5,000 cells per well in a six-well plate. After 48 hours, cells were fed with mTeSR1 (without Bleb) every 24 hours until the next passage. To minimize cell stress, no antibiotics were used in RPMI (Gibco, USA) and supplement media (10% fetal bovine serum (FBS), 2.5% penicillin). Cells were maintained at 37°C and 5% $CO_2$ and passaged every 3-4 days at ~1 x $10^{\wedge 5}$ – 2 x $10^{\wedge 6}$ cells/ml in uncoated flasks. Cells were routinely tested for mycoplasma using MycoAlert (Lonza, Switzerland, No. LT07).

***Retinal organoid culturing***

H9 CRX:tdtomato$^{+}$ hESCs were dissociated in Accutase at 37°C for 12 min and seeded in 50 µl of mTeSR1 at 3,000 cells/well into 96-well ultra-low adhesion round bottom Lipidure coated plates (AMSBIO, MA, USA, No.51011610). Cells were placed in hypoxic conditions (10% $CO_2$ and 5% $O_2$) for 24 hours to enhance survival. Cells naturally aggregated by gravity over 24 hours. On day 1, cells were moved to normoxic conditions (5% $CO_2$). On days 1- 3, 50 µl of BE6.2 media, supplementary **Table S1**) containing 3 µM Wnt inhibitor (IWR1e, EMD Millipore, MA, USA, No. 681669,) and 1% (v/v) Matrigel were added to each well. On days 4-9, 100 µl of media were removed from each well, and 100 µl of media were added. On days 4-5, BE6.2 media containing 3 µM Wnt inhibitor and 1% Matrigel was added. On days 6-7, BE6.2 media containing 1% Matrigel was added. On days 8-9, BE6.2 media containing 1% Matrigel and 100 nM Smoothened agonist (SAG, EMD Millipore, No. 566660) was added. On day 10, aggregates were transferred to 15 mL tubes, rinsed 3X in DMEM (Gibco, No. 11885084), and resuspended in BE6.2 with 100 nM SAG in untreated 10 cm polystyrene petri dishes. From this point on, media was changed every other day. Aggregates were monitored and manually separated if stuck together or to the bottom of the plate. On day 11, retinal vesicles were manually dissected using sharpened tungsten needles. After dissection, cells were transferred into 15 mL tubes and washed 2X with 5 mLs of DMEM. On days 14-17, long-term retina (LTR, supplementary **Table S2**) media with 100 nM SAG was added. On days 18-21, cells were maintained in LTR and washed 2X with 5 mLs of DMEM, before being transferred to new plates to wash off dead cells. To increase survival and differentiation, 1 µM all-trans retinoic acid (ATRA; R2625; Sigma) was added to LTR medium from days 22-138. 10 µM Gammasecretase inhibitor (DAPT, EMD Millipore, No. 565770) was added to LTR from days 28-42. Retinal organoids were grown at low density (10-20 per 10 cm dish) to reduce aggregation.

***Animals***

All animal experiments were carried out in accordance with the ARVO Statement for the Use of Animals in Ophthalmic and Vision Research. All procedures were approved by the Johns Hopkins University Animal Care and Use Committee (approval M016M17). The C3H/HeJ-*Pde6*$^{Rd1/Rd1}$ (referred to as *Rd1*), and NOD.Cg-*Prkdc*$^{scid}$/J (referred to as *NOD/Scid*) mice of either gender (aged 6 to 8 weeks) were obtained from Jackson Laboratory (Bar Harbor, ME, USA). All mice were housed in cages under a 12:12-hour light-dark cycle with water and food provided *ad libitum*.

***Recipient mice***

We created a recipient mouse model with immune-deficiency and retinal degeneration (referred to as *Rd1/NS*) by crossbreeding *Rd1* mice and *NOD/Scid* mice (aged 8 weeks). The breeding strategy was performed as previously reported [1]. Genomic DNA of the third-generation offspring was extracted from ear biopsies and genotyped by Transnetyx Tag Center (Cordova,TN, USA). Primers were listed in supplementary **Table S3**. Eyes of adult *Rd1/NS* mice (n=3) were collected to characterize photoreceptor degeneration using immunohistochemistry (IHC) staining, as previously reported [2]. Flow cytometry was performed using spleen biopsies of adult *Rd1/NS* mice (n=3) to confirm the deficiency of T cells and B cells, as previously described [3]. Phenotyping data of *Rd1/NS* mice are shown in supplementary **Fig. S1**.

***Preparation of donor cells***

Donor retinal organoid cells (harvested as micro-dissected multilayered retinal fragments) were obtained from CRX:tdTomato$^+$ hESC-derived retinal organoids (aged 134 days, n=4). The cultured human retinal organoids were imaged using a fluorescent microscope (Carl Zeiss, Jena, Germany). The images were used as a reference to isolate the CRX:tdTomato$^+$ cluster from the donor retinal organoids. The isolated CRX:tdTomato$^+$ retinal organoids clusters were then cut into $1 \times 1$ mm$^{\wedge 2}$

or $1 \times 2\ mm^{\wedge 2}$ microdissected fragments using a 27-gauge horizontal curved scissors (VitreQ, Kingston, NH, USA) under a dissection microscope. Donor cells were transplanted within two hours of isolation.

***Transplantation of donor cells***

Donor retinal organoid fragments were transplanted into the subretinal space of *Rd1/NS* mice (aged 6 to 8 weeks, n=16 eyes), as previously reported [4]. Briefly, recipient mice were anesthetized by intraperitoneal injection of ketamine (100 mg/kg body weight) and xylazine hydrochloride (20 mg/kg body weight). Mouse pupils were dilated with 1% (wt/vol) tropicamide (Bausch & Lomb, Rochester, NY, USA). Mouse corneas were covered with Sodium Hyaluronate (Healon GV, Abbott Medical Optics Inc. CA, USA) and cover glasses (Deckglaser, USA) to facilitate transpupillary visualization. The donor retinal organoid fragments were loaded into the bevel of a 26G microneedle with the photoreceptor side facing down, gently aspirated into the attached micro-syringe (Hamilton, Reno, NV, USA), then tangentially injected into the subretinal space through the sclera of the recipient mice. Successful injection was verified by direct visualization through the dilated pupil of the recipient under the surgical microscope (Leica, Wetzlar, Germany).

***Single cell RNA sequencing***

Single cell RNA sequencing (scRNA-seq) was performed on dissociated cells from transplanted and cultured retinal organoids using the Chromium platform (10X Genomics). Briefly, retinal organoid cells were dissociated into a single cell suspension using the Papain Dissociation System (Worthington) for 60 minutes at 37°C, with gentle mixing every 5 minutes, before stopping the reaction using ovomucoid protease inhibitors. Cells were centrifuged and resuspended in ice-cold PBS containing 0.04% bovine serum albumin (BSA) and 0.5 U/µl RNase inhibitor and were filtered through a 40-µm Flowmi cell strainer (Bel-Art SP). Cell counts and viability were assessed by Trypan blue staining before loading 6000 cells on a Chromium Single Cell system using Next GEM 3' reagent v3.1 kits. Libraries were pooled and sequenced on Illumina NextSeq 500 with ~50,000 reads per cell. The Cell Ranger 4 (10X Genomics) pipeline was used to process the raw sequencing reads for demultiplexing, alignment to the GRCh38 human reference genome and generating the cell-by-gene count matrix for downstream analysis. The generated cell-by-gene count matrices were analyzed using the Seurat ver3 R package [5]. We filtered out cells that had UMIs less than 300 or greater than 50000 and with a mitochondrial fraction of greater than 20%. Doublets were identified and removed using the DoubletFinder R package [6]. Log-normalization, scaling, UMAP dimensional reduction and clustering were performed using the standard Seurat pipeline. Quality control of scRNA-seq data were shown in supplementary **Fig. S5**. Major retinal cell types were identified using previously identified cell type markers [7]. Enriched genes from the brain/spinal-like cell cluster were compared to the ASCOT [8] gene expression summaries of public RNA-Seq data to determine its classification. Differential gene tests were performed by Seurat's *FindMarkers* function using the Wilcoxon rank sum test with default parameters [5]. Hierarchical clustering was used to group the differentially expressed genes. The UCell R package [9] was used to calculate the migration potential score or the proliferation score (supplementary **data files S1, S2**). The gene sets were constructed by identifying enriched genes within the gene ontology terms cell migration and cell motility for the migration potential score and cell division for the proliferation score respectively. The Seurat integration functions (*SelectIntegrationFeatures*, *FindIntegrationAnchors* and *IntegrateData*) were used to integrate the organoid data onto the human retinal developmental dataset [10]. Monocle 3 [11] was used to perform pseudotime analysis and identify trajectory routes within the data.

***Histology***

Four and a half months post-transplantation, the recipient mice were sacrificed with over-dose anesthesia and pre-fixed by heart-perfusion with 4% paraformaldehyde (PFA) (Electron Microscopy Sciences, Hatfield, PA, USA) in PBS. Eyes were gently removed, post-fixed in 4% PFA/PBS for one hour at room temperature (RT), and dehydrated in a sucrose gradient (10%, 20%, 30%), then blocked in optimal cutting temperature compound (Sakura Finetek, Torrance, CA, USA). Cultured retinal organoids were fixed in 4% PFA at RT for 15 minutes (min), dehydrated in gradient sucrose (10%, 20%, 30%), and blocked in the OCT compound. OCT-blocked recipient mouse eyes and cultured retinal organoids were cut into 7-10 μm thick cryosections using a microtome (CM 1850; Leica) for histological staining.

RNAscope and IHC counter-staining was performed according to the manufacturer's protocol (Advanced Cell Diagnostics (ACD), see Protocol #MK 51-150, Appendix D.). Briefly, cryosections of recipient mice eyes and cultured retinal organoids were rinsed with PBS, baked in a HybEZTM oven (ACD, USA) for 30 min at 60°C, and post-fixed in pre-chilled 4% PFA in PBS for 15 min at 4°C. Slides were dehydrated in gradient ethanol (50%, 70%, 100%), treated with hydrogen peroxide (10 min at RT), then subjected to target retrieval using the Co-detection Target Retrieval solution (ACD, Cat. No. 323180) at 98-102°C for 5 min. After rinsing in distilled water (2 min x 2) and PBS-T (5 min x 1), the slides were incubated with diluted primary antibody at 4°C overnight. On day 2, slides were post-fixed with 4% PFA for 30 min at RT, treated with protease III at 40°C for 30min, and subjected to RNAscope staining using the RNAscope Multiplex Fluorescent V2 assay according to the manufacturer's protocol (ACD, RNAscope USM-323100, see "fixed-frozen tissue sample protocol"). Briefly, RNA probe hybridization was performed with the HybEZTM oven for two hours at 40°C. Slides were then assigned for three series of amplification, fluorochromes combination, and HRP blocking. After the RNAscope procedure, slides were incubated with secondary antibody at RT for one hour, counter stained with DAPI, and mounted with Prolong Diamond (Life Technology, Carlsbad, CA, USA). The RNA probes, fluorophores, primary antibodies, and secondary antibodies used were listed in supplementary **Table S4**. Negative and positive multiplex control probes staining were run in parallel with the target probes following the same protocol (data shown in supplementary **Fig. S6**).

IHC staining was performed as previously described [4]. Briefly, cryosections of transplanted *Rd1/NS* mice and cultured retinal organoids were rinsed with PBS (5 min x 1), permeabilized, and blocked with a mixture of 0.1% Triton-X100 and 5% goat serum in PBS for one hour at RT. The slides were rinsed in PBS (5 min x 3), incubated with primary antibodies at 4°C overnight, incubated with secondary antibodies at RT for one hour, then counter stained with DAPI and mounted using ProLong Diamond mounting media. The primary antibodies and secondary antibodies used were listed in supplementary **Table S5**.

***Quantification of donor cell migration of recipient retina***

For migratory distance quantification of transplanted retinal organoid cells, retinal sections from recipient mice were stained with human nuclear specific antibodies HNA (Sigma-Aldrich, MO, USA) or Ku80 (Thermo Fisher Scientific, MA, USA). Tile scan images were collected using Confocal LSM 880 (Zeiss, Oberkochen, Germany) for distance quantification. The migratory distance of transplanted retinal organoids was defined as the shortest distance between the migratory cells and the nearest graft edge (i.e., the graft-left migratory cells to the left endpoint of the graft; the graft-right migratory cells to the right endpoint of the graft). We used a mathematical method to facilitate distance quantification. Specifically, the graft edge was defined as a "starting point" and the migratory cells in different retinal laminae (RGC, IPL, INL, RPE/C) were manually targeted, both processed with the "Cell Counter" plugin in ImageJ. The cell coordinates were automatically collected to quantify the X and Y axial distances of individual cells by the Cell Counter. The axial distance of the graft edge (starting point) was referred to as "$X_{start}$" and "$Y_{start}$". The axial distance of the migratory cells was referred to as "$X_{migratory}$" and "$Y_{migratory}$". The migratory distance was computed in R platform (see supplementary **coding file S1**) following the formula:

$$Migrating\ distance\ = \sqrt{(X_{start} - X_{migratory})^2\ +\ (Y_{start} - Y_{migratory})^2}$$

The unit of the migrating distance was converted from pixel to micron according to the image scale.

For cell quantification, the number of positively stained cells was manually counted using the "Cell Counter" plugin in ImageJ. The representative pre-synapse graphs of the transplanted and cultured retinal organoids were drawn by Imaris software (Version 9.5.0, Bitplane AG, Zurich, Switzerland).

***Electrophysiology***

The electrophysiological recording was performed on the transplanted photoreceptors eight months post-transplantation to measure their physiological properties. We were able to test only one recipient mouse (the second recipient mouse died before the assay during the long-term observation). The recipient's eyes were gently pulled out from the recipient mouse and put in Ames' medium (Sigma No. A1420). Retinas with transplanted retinal organoids were dissected by removing the corneas and lens under infrared light, attached to a piece of filter, sectioned into 200μm slices, and transferred to a recording chamber. The CRX:tdTomato$^+$ photoreceptors of the transplanted retinal organoids were targeted under an epifluorescence microscope for consequent whole-cell patch-clamp recording. Fluorescent signal was imaged by a Nikon CCD camera with data acquisition synchronized with a 20-ms flash of epi-fluorescence excitation light. The total exposure time to excitation light before recording was <500 ms. During recording, retina was perfused with Ames' medium bubbled with 95% $O_2$/5% $CO_2$. Patch electrodes (5-7 MΩ) were pulled from borosilicate capillaries (GC150-10, Harvard Apparatus) and filled with an internal solution containing typically (in mM): 120 K-gluconate, 5 NaCl, 4 KCl, 10 HEPES, 2 EGTA, 4 ATP-Mg, 0.3 GTP-$Na_2$, and 7 Phosphocreatine-Tris, with pH adjusted to 7.3 with KOH. Whole-cell patch-clamp recording was made at 30–32$^{o}$C with an Axon Instruments Multiclamp 700B amplifier. Series resistance of patch electrodes was 10–30 MΩ. Liquid-junction potential (measured to be -13 mV) has been corrected. In voltage-clamp mode, recorded cells were held at -40 mV, followed by voltage steps of 100-ms (-70 mV to -10 mV). All procedures were carried out in the darkroom to avoid photoreceptor bleaching.

**Supplementary Figure 1**

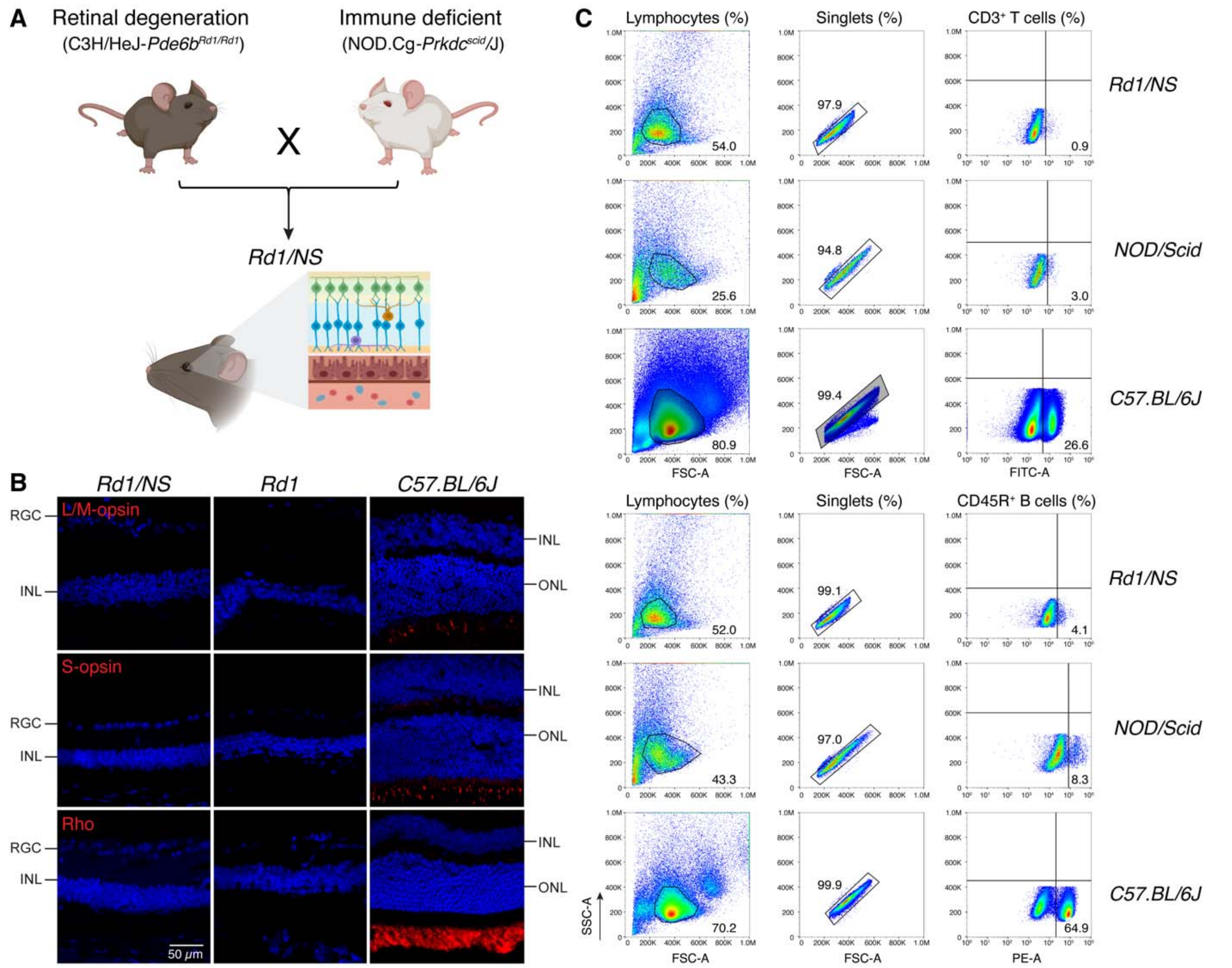


**Fig. S1. Breeding and phenotyping of the recipient *Rd1/NS* mice. (A)** Schematic showed the recipient *Rd1/NS* mice were generated by crossbreeding C3H/HeJ-*Pde6b*$^{Rd/Rd11}$ (*Rd1)* and NOD.Cg-*Prkdc*$^{scid}$/J (*NOD/Scid*) mice. **(B)** IHC staining showed fully degenerated photoreceptor cells and negative expression of L/M-opsin, S-opsin, and Rhodopsin (Rho) in adult *Rd1/NS* mice and *Rd1* mice. *C57.BL/6J* mice served as *wild-type* controls. **(C)** Flow cytometry analysis showed the deficiency of CD3$^+$ T cells and CD45R$^+$ B cells in *Rd1/NS* mice, corresponding to the immune deficient phenotype of *NOD/Scid* mice. *C57.BL/6J* mice served as *wild-type* controls.

**Supplementary Figure 2**

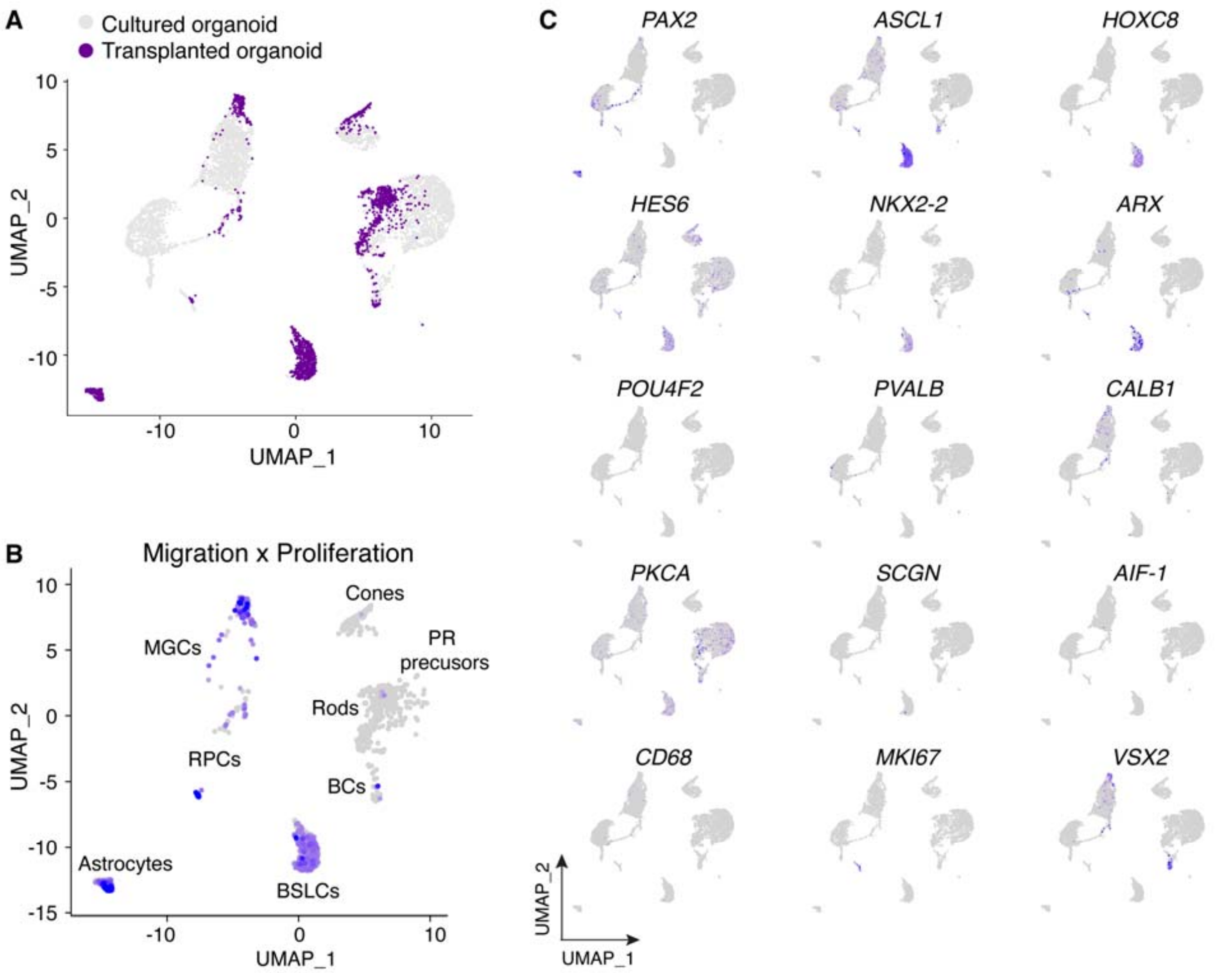


**Fig. S2. UMAP plots of migration and proliferation cell clusters. (A)** UMAP plot colored cell clusters of cultured (grey) and transplanted retinal organoids (purple). **(B)** UMAP plot colored cell clusters sharing transcriptomic characteristics of migration and proliferation. **(C)** UMAP plots displayed the expression of marker genes in cell clusters of transplanted and cultured retinal organoids.

**Supplementary Figure 3**

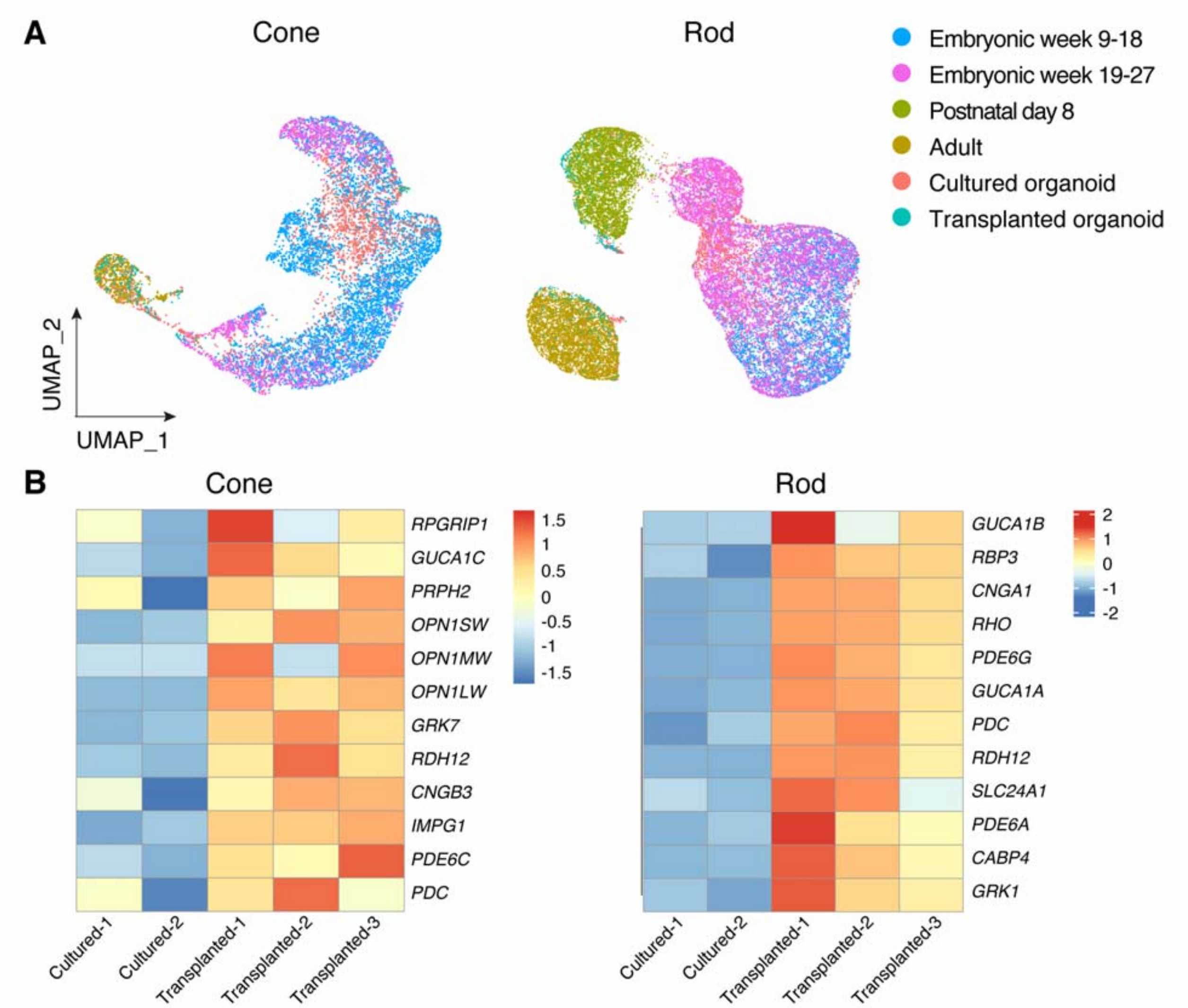


**Fig. S3. Upregulation of cone and rod marker genes in transplanted retinal organoids. (A)** UMAP plots showed cells colored by sample libraries, including human retina developmental datasets (cone: n = 7,654 cells, rod: n = 25,186 cells), cultured retinal organoids (cone: n =1,639 cells, rod: n=1,469 cells), and transplanted retinal organoids (cone: n=210 cells, rod: n =504 cells). **(B)** Heatmaps demonstrated the upregulation of marker genes specific for cone and rod photoreceptors in transplanted retinal organoids (including three independent replicates "Transplanted-1, Transplanted-2, Transplanted-3"), compared to cultured retinal organoids (including two independent replicates "Cultured-1, Cultured-2").

**Supplementary Figure 4**

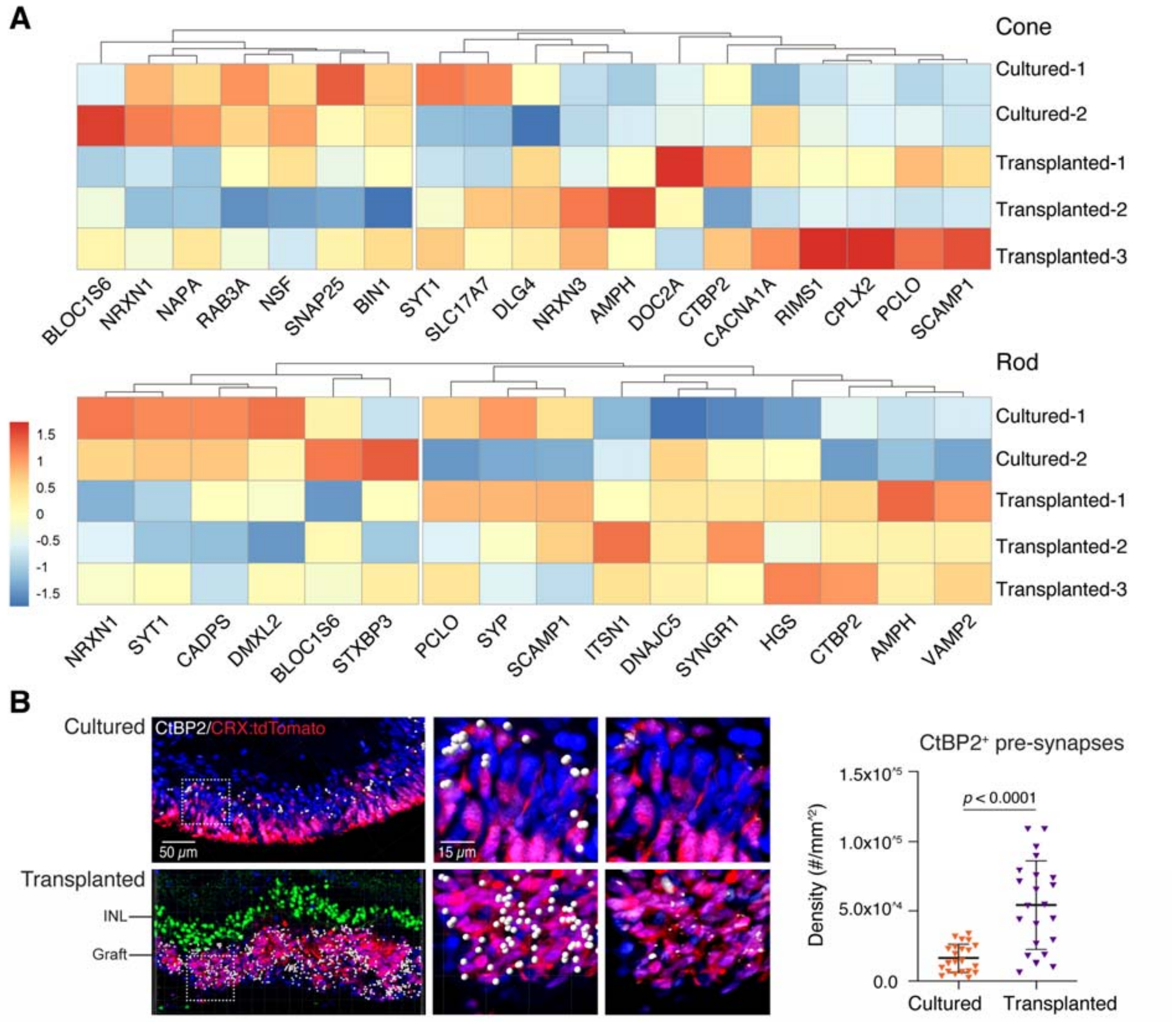


**Fig. S4. Identification and quantitation of pre-synaptic markers in cultured and transplanted retinal organoids. (A)** The heatmaps demonstrate that several synapse-related genes were upregulated in the transplanted retinal organoids (including cones and rods) compared to the cultured retinal organoids. **(B)** IHC staining and quantification demonstrated that $CtBP2^{+}$ synaptic ribbons in photoreceptors ($CRX:tdTomato^{+}$) were significantly more abundant in transplanted than cultured retinal organoids. IHC staining of SCGN (green) demonstrated the recipient bipolar layer. The anti-human nuclear antibody (HNA, blue) was used to label human cells. *Abbreviation: INL: inner nuclear layer.*

**Supplementary Figure 5**

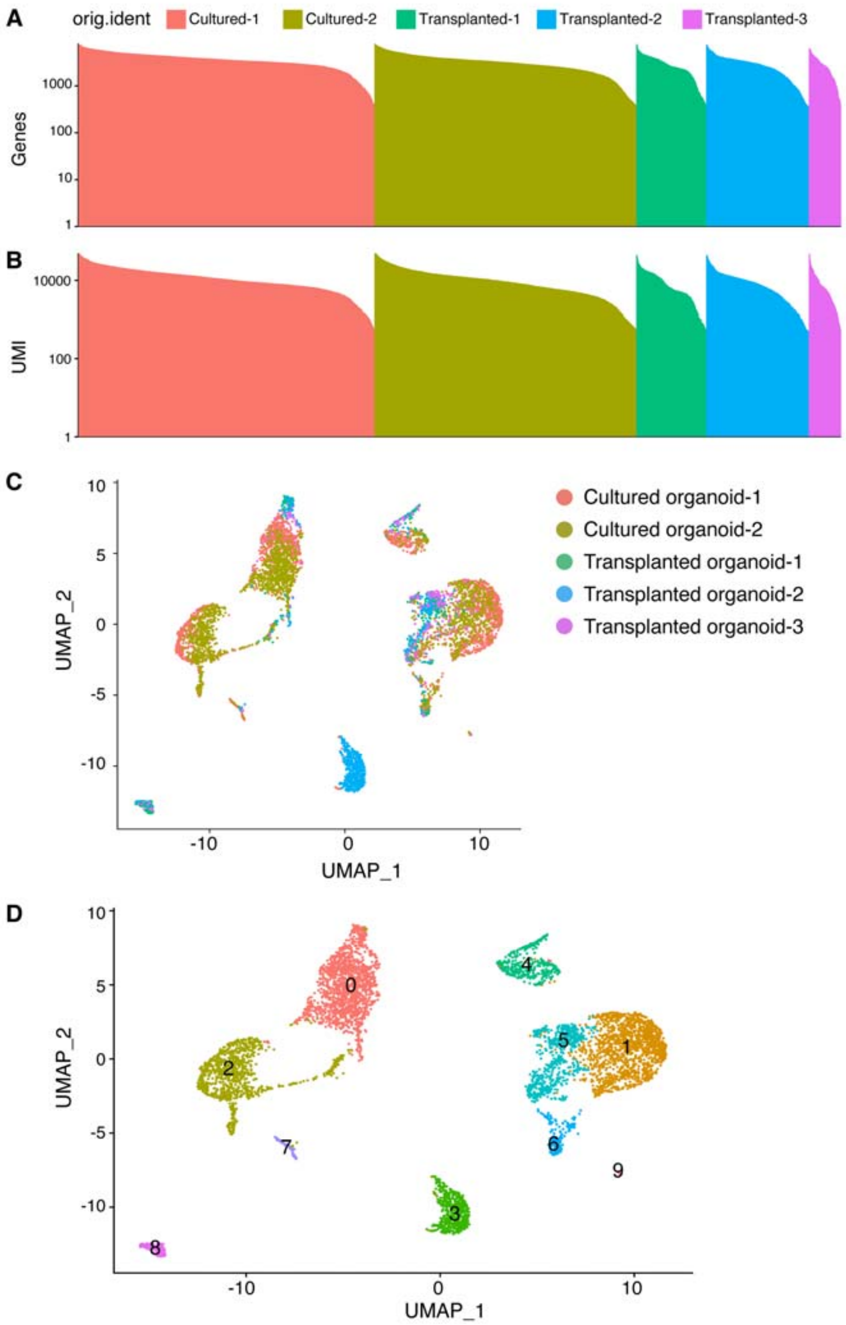


**Fig. S5. Quality control of scRNA-seq data. (A)** Number of genes and **(B)** unique molecular identifiers (UMI) per cell. Each bar is a cell and is colored by the sample library and ordered along the x-axis in descending order. **(C)** UMAP plot showing cells colored by sample library. **(D)** UMAP plot showing 10 (0-9) transcriptionally distinct cell clusters.

**Supplementary Figure 6**

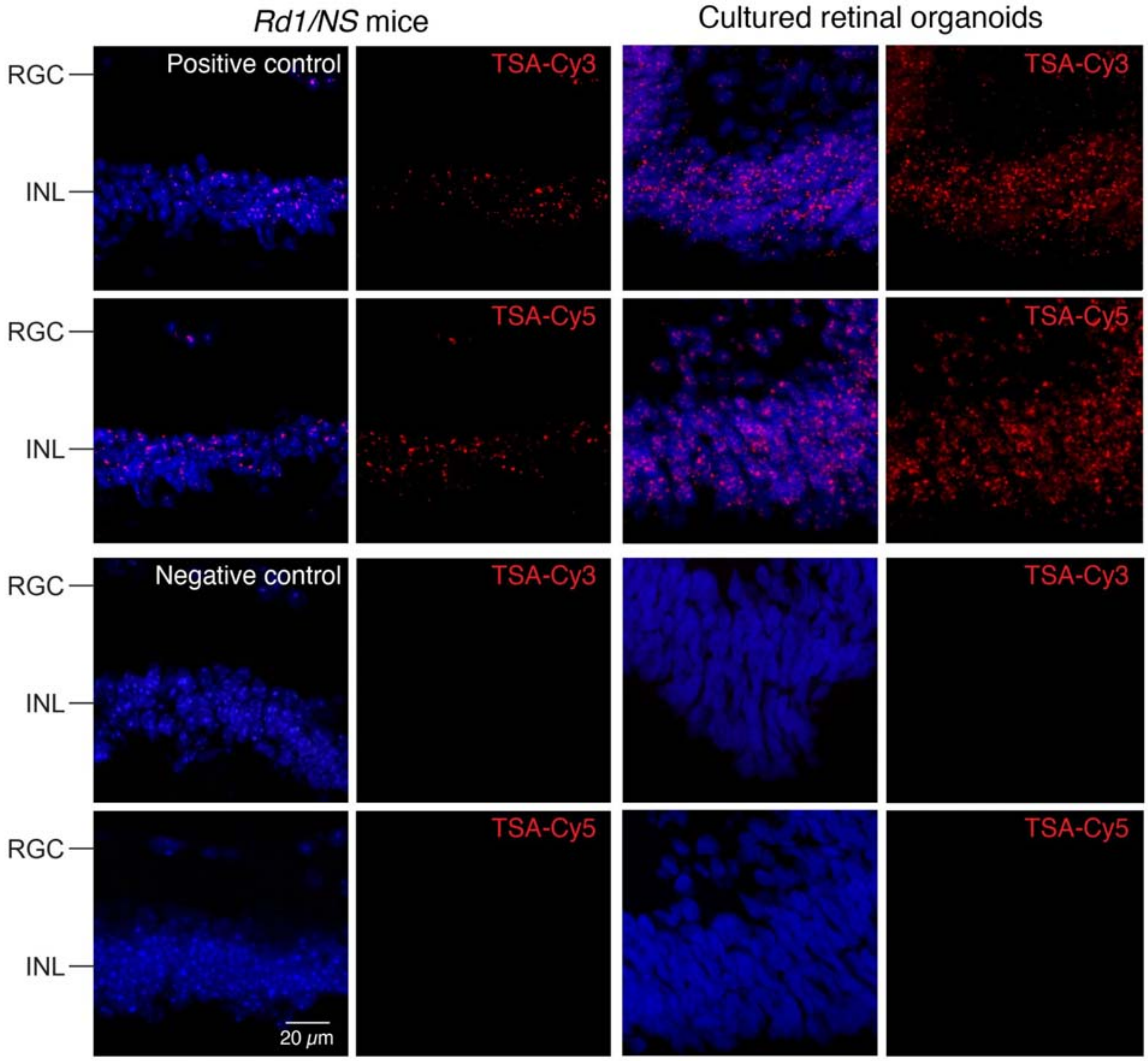


**Fig. S6. RNAscope staining of positive and negative control probes.** Cryosections of non-transplanted *Rd1/NS* mice and cultured retinal organoids were stained with 3-plex positive and negative control probes in combination with TSA-Cy3 or TSA-Cy5 fluorophores. Positive probes target common housekeeping genes *PPIB* (Cy3) and *POLR2A* (Cy5). Negative probe targets the bacterial dapB gene.

**Table S1. BE6.2 media for early retinal differentiation.**

| Reagent | Concentration | Source | Catalog Number |
|---|---|---|---|
| DMEM | — | Gibco | 11885084 |
| Minus vitamin A | 2% | Gibco | 12587010 |
| Glutamax | 1% | Gibco | 35050061 |
| NEAA | 1% | Gibco | 11140050 |
| Sodium pyruvate | 1mM | Gibco | 11360070 |
| NaCl | 0.87 mg/mL | Sigma-Aldrich | S9888 |
| **E6 supplement** | **2.5%** | | |
| Insulin | 970 ug/mL | Roche | 11376497001 |
| Holo-transferrin | 535 ug/mL | Sigma-Aldrich | T0665 |
| L-ascorbic acid | 3.20 mg/mL | Sigma-Aldrich | A8960 |
| Sodium selenite | 0.7 ug/mL | Sigma-Aldrich | S5261 |

**Table S2. Long-term retina media (LTR) for retinal organoids culturing.**

| Reagent | Concentration | Source | Catalog Number |
|---|---|---|---|
| DMEM | — | Gibco | 11885084 |
| F12 | 25% | Gibco | 11765062 |
| B27 | 2% | Gibco | 17504044 |
| NEAA | 1% | Gibco | 11140050 |
| Fetal bovine serum | 10% | Gibco | 16140071 |
| Sodium pyruvate | 1mM | Gibco | 11360070 |
| Glutamax | 1% | Gibco | 35050061 |
| Taurine | 1 mM | Sigma-Aldrich | T-8691 |

**Table S3. Forward and reverse primer sequences used for mice genotyping.**

| Gene | F primer (5’ to 3’) | R primer (5’ to 3’) |
|---|---|---|
| *Rd1 Wild type* | ACTCTGTGGCCTCAAAGATACATC | TGCAGGTCACAGAATCATCATAACA |
| *Rd1 Mutant* | GGGTCTCCTCAGATTGATTGACTAC | GTCACTCTGTGGCCTCAAAGAT |
| *NOD/Scid* | TGTAACGGAAAAGAATTGGTATCCACA | GTTGGCCCCTGCTAACTTTCT |

**Table S4. Reagents used for RNAscope and immunohistochemistry counterstaining.**

| Reagents | Dilution | Source | Catalog Number |
|---|---|---|---|
| **RNA probes** | | | |
| PAX2-Hs | No dilution | Advanced Cell Diagnostics | 442541 |
| HES6-Hs | 1:50 | Advanced Cell Diagnostics | 521301-C2 |
| ASCL1-Hs | 1:50 | Advanced Cell Diagnostics | 459721-C2 |
| NKX2-2-Hs | No dilution | Advanced Cell Diagnostics | 821401 |
| HOXC8-Hs | No dilution | Advanced Cell Diagnostics | 506531 |
| ARX-Hs | 1:50 | Advanced Cell Diagnostics | 486711-C2 |
| VSX2-Hs | 1:50 | Advanced Cell Diagnostics | 493031-C2 |
| 3-plex positive control-Mm | No dilution | Advanced Cell Diagnostics | 320881 |
| 3-plex positive control-Hs | No dilution | Advanced Cell Diagnostics | 320861 |
| 3-plex negative control | No dilution | Advanced Cell Diagnostics | 320871 |
| **TSA-fluorophores** | | | |
| TSA plus-Cy3 | 1:1500 | AKOYA Biosciences | NEL744001KT |
| TSA plus-Cy5 | 1:1500 | AKOYA Biosciences | NEL745001KT |
| **Primary antibodies** | | | |
| Sheep anti-Ki67 | 1:40 | R&D Systems | AF7617 |
| Rabbit anti-Ku80 | 1:50 | Thermo Fisher Scientific | MA5-32212 |
| **Secondary antibodies** | | | |
| Donkey anti-Sheep 647 | 1:200 | Thermo Fisher Scientific | A-21448 |
| Donkey anti-Rabbit 488 | 1:200 | Thermo Fisher Scientific | A21206 |

**Table S5. Antibodies used for immunohistochemistry staining.**

| Reagents | Dilution | Source | Catalog Number |
|---|---|---|---|
| **Primary antibodies** | | | |
| Rabbit anti-Brn3b | 1:100 | Abcam | Ab56026 |
| Goat anti-Parvalbumin | 1:200 | LifeSpan bioSciences | LS-B3601 |
| Mouse anti-Calbindin | 1:500 | Sigma-Aldrich | C9848 |
| Goat anti-PKCα | 1:200 | R&D Systems | AF5340 |
| Goat anti-SCGN | 1:200 | Thermo Fisher Scientific | PA5-47664 |
| Rabbit anti-IBA1 | 1:250 | Abcam | Ab178680 |
| Rabbit anti-CD68 | 1:100 | Abcam | Ab125212 |
| Rabbit anti-Ki67 | 1:500 | Thermo Fisher Scientific | MA5-14520 |
| Rabbit anti-L/M-opsin | 1:500 | Kerafast | EDK101 |
| Rabbit anti-S-opsin | 1:500 | MilliporeSigma | Ab5407 |
| Rabbit anti-Rhodopsin | 1:500 | Abcam | Ab3424 |
| Mouse anti-human nuclear antibody (HNA) | 1:1000 | MilliporeSigma | Mab1281 |
| **Secondary antibodies** | | | |
| Goat anti-Rabbit 488 | 1:500 | Abcam | Ab150077 |
| Goat anti-Mouse 488 | 1:500 | Thermo Fisher Scientific | A-11001 |
| Goat anti-Mouse 647 | 1:500 | Thermo Fisher Scientific | A32728 |
| Donkey anti-Goat 488 | 1:500 | Abcam | Ab150129 |
