## Supplemental code for "Transretinal migration of astrocytes and brain/spinal cord-like cells arising from transplanted human retinal organoids"

### R coding algorithms for migrating distance quantification

```
library(data.table)
library(plyr)

################################################################################
#
# Extract the X,Y coordinate of migrated cells, and generate subsequent distance data presented in Table 1
#
################################################################################

#---> Extracting data exported from ImageJ

common_path = "~/Desktop/cell invasion/Processed_Image"

files_to_read = list.files(
  path = common_path,       # directory to search within
  pattern = ".*(Rd1-NS).*csv$", # regex pattern
  recursive = TRUE,         # search subdirectories
  full.names = TRUE         # return the full path
)

#---> Hypothetical Researcher has 37 retina slides to quantify
#     and wants to localize 5 cell types per retina slides. So that's
#     37 rows and 6 columns (including cell id) in the data list

data_lst = lapply(files_to_read, read.csv)  # read all the matching files
celltype_summary = data.frame(matrix(ncol = 6, nrow = 37))
colnames (celltype_summary) <- c("File_name", "Type_1", "Type_2", "Type_3", "Type_4", "Type_5")

#---> Calculating the distance from starting point for each cell
#     categorized by cell types

for (i in 1:length(data_lst)){
  data_lst[[i]]$Address <- rep(files_to_read[i],nrow(data_lst[[i]]))
  File_Name <-  files_to_read[i]
  Type_1 <- sum(data_lst[[i]]$Type == 1)
  Type_2 <- sum(data_lst[[i]]$Type == 2)
  Type_3 <- sum(data_lst[[i]]$Type == 3)
  Type_4 <- sum(data_lst[[i]]$Type == 4)
```

```
  Type_5 <- sum(data_lst[[i]]$Type == 5)
  celltype_summary[i,1:6] = c(File_Name, Type_1, Type_2, Type_3, Type_4, Type_5)
  print(c(File_Name, Type_1, Type_2, Type_3, Type_4, Type_5))
  for (j in 1:nrow(data_lst[[i]])){
  data_lst[[i]][j,"Z.µm."]=sqrt(( data_lst[[i]][j,"X.µm."]- data_lst[[i]][1,"X.µm."])^2 +
                                ( data_lst[[i]][j,"Y.µm."]- data_lst[[i]][1,"Y.µm."])^2) #distance calculation
  }
}

#---> Exporting analyzed invasion distance summary

dat1<-ldply(data_lst)

write.table(as.data.frame(dat1),file="Detialed_Result.csv", quote=F,sep=",",row.names=F)
write.table(as.data.frame(celltype_summary),file="Celltype_Summary.csv",
quote=F,sep=",",row.names=F)
```
